## Supplementary material for "Small Training Dataset Convolutional Neural Networks for Application Specific Super-Resolution Microscopy": super-resolution latex file: ReadMe.txt

SPIE e-journal LaTeX style files
----------------------------------
Updated: 2015/07/24

Basic LaTeX files:
spieman.cls,  class file to format manuscript
spiejour.bst  BibTeX bibliography style file to format reference list

Demonstration document including formatting instructions:
article.tex, LaTeX source file
report.bib, BibTeX bibliography file used by article.tex
fig2.eps, EPS image used by article.tex combining two images
mcr3b.eps, EPS image used by article.tex
satellite.eps, EPS image used by article.tex
  
Example of the final formatted manuscript obtained using the above style files:
article.pdf, in Adobe Portable Document Format
