## Supplementary material for "Small Training Dataset Convolutional Neural Networks for Application Specific Super-Resolution Microscopy": super-resolution latex file: report.bib

%% report.bib - contains BiBtex input for bibliography
@article{zhang2017beyond,
  title={Beyond a {G}aussian denoiser: Residual learning of deep {CNN} for image denoising},
  author={Zhang, Kai and Zuo, Wangmeng and Chen, Yunjin and Meng, Deyu and Zhang, Lei},
  journal={IEEE Transactions on Image Processing (TIP)},
  volume={26},
  number={7},
  pages={3142--3155},
  year={2017},
  publisher={IEEE}
}

%ML
@book{goodfellow2016deep,
  title={Deep {L}earning},
  author={Goodfellow, Ian and Bengio, Yoshua and Courville, Aaron and Bengio, Yoshua},
  volume={1},
  number={2},
  year={2016},
  publisher={MIT press Cambridge}
}

%new citations
@article{lichtman2005fluorescence,
  title={Fluorescence {M}icroscopy},
  author={Lichtman, Jeff W and Conchello, Jos{\'e}-Angel},
  journal={Nature Methods},
  volume={2},
  number={12},
  pages={910--919},
  year={2005},
  publisher={Nature Publishing Group}
}

%spie paper DL
@inproceedings{mannam2021deep,
  title={Deep learning-based super-resolution fluorescence microscopy on small datasets},
  author={Mannam, Varun and Zhang, Yide and Yuan, Xiaotong and Howard, Scott},
  booktitle={Single Molecule Spectroscopy and Superresolution Imaging XIV},
  volume={11650},
  pages={116500O},
  year={2021},
  organization={International Society for Optics and Photonics (SPIE Photonics West)}
}


%from XY super-resolution base papers:
@article{STED,
  title={Breaking the diffraction resolution limit by stimulated emission: stimulated-emission-depletion fluorescence microscopy},
  author={Hell, Stefan W and Wichmann, Jan},
  journal={Optics {L}etters.},
  volume={19},
  number={11},
  pages={780--782},
  year={1994},
  publisher={Optical Society of America}
}

@article{STORM,
  title={Sub-diffraction-limit imaging by stochastic optical reconstruction microscopy ({STORM})},
  author={Rust, Michael J and Bates, Mark and Zhuang, Xiaowei},
  journal={Nature Methods},
  volume={3},
  number={10},
  pages={793},
  year={2006},
  publisher={Nature Publishing Group}
}

@article{PALM,
  title={Imaging intracellular fluorescent proteins at nanometer resolution},
  author={Betzig, Eric and Patterson, George H and Sougrat, Rachid and Lindwasser, O Wolf and Olenych, Scott and Bonifacino, Juan S and Davidson, Michael W and Lippincott-Schwartz, Jennifer and Hess, Harald F},
  journal={Science},
  volume={313},
  number={5793},
  pages={1642--1645},
  year={2006},
  publisher={American Association for the Advancement of Science}
}

@article{SIM,
  title={Surpassing the lateral resolution limit by a factor of two using structured illumination microscopy},
  author={Gustafsson, Mats GL},
  journal={Journal of Microscopy},
  volume={198},
  number={2},
  pages={82--87},
  year={2000},
  publisher={Wiley Online Library}
}

@article{SIM_PNAS,
  title={Nonlinear structured-illumination microscopy: wide-field fluorescence imaging with theoretically unlimited resolution},
  author={Gustafsson, Mats GL},
  journal={Proceedings of the National Academy of Sciences (PNAS)},
  volume={102},
  number={37},
  pages={13081--13086},
  year={2005},
  publisher={National Acad Sciences}
}

@article{NSIM,
  title={Nonlinear structured-illumination microscopy with a photoswitchable protein reveals cellular structures at 50-nm resolution},
  author={Rego, E Hesper and Shao, Lin and Macklin, John J and Winoto, Lukman and Johansson, G{\"o}ran A and Kamps-Hughes, Nicholas and Davidson, Michael W and Gustafsson, Mats GL},
  journal={Proceedings of the National Academy of Sciences (PNAS)},
  volume={109},
  number={3},
  pages={E135--E143},
  year={2012},
  publisher={National Acad Sciences}
}

@article{GSOS,
  title={Generalized stepwise optical saturation enables super-resolution fluorescence lifetime imaging microscopy},
  author={Zhang, Yide and Benirschke, David and Abdalsalam, Ola and Howard, Scott S},
  journal={Biomedical Optics Express},
  volume={9},
  number={9},
  pages={4077--4093},
  year={2018},
  publisher={Optical Society of America}
}

%from conference import
%cvpr for noise2noise model
@inproceedings{zhang2019poisson,
  title={A poisson-gaussian denoising dataset with real fluorescence microscopy images},
  author={Zhang, Yide and Zhu, Yinhao and Nichols, Evan and Wang, Qingfei and Zhang, Siyuan and Smith, Cody and Howard, Scott},
  booktitle={Proceedings of the IEEE Conference on Computer Vision and Pattern Recognition},
  pages={11710--11718},
  year={2019}
}

%vm denoising
@inproceedings{mannam2020instant,
  title={Instant Image Denoising Plugin for {ImageJ} using Convolutional Neural Networks},
  author={Mannam, Varun and Zhang, Yide and Zhu, Yinhao and Howard, Scott},
  booktitle={Microscopy Histopathology and Analytics},
  pages={MW2A--3},
  year={2020},
  organization={Optical Society of America}
}


%vm: review
@article{mannam2020machine,
  title={Machine learning for faster and smarter fluorescence lifetime imaging microscopy},
  author={Mannam, Varun and Zhang, Yide and Yuan, Xiaotong and Ravasio, Cara and Howard, Scott S},
  journal={Journal of Physics: Photonics},
  volume={2},
  number={4},
  pages={042005},
  year={2020},
  publisher={IOP Publishing}
}
@inproceedings{zhang2018residual,
  title={Residual dense network for image super-resolution},
  author={Zhang, Yulun and Tian, Yapeng and Kong, Yu and Zhong, Bineng and Fu, Yun},
  booktitle={Proceedings of the IEEE Conference on Computer Vision and Pattern Recognition (CVPR)},
  pages={2472--2481},
  year={2018}
}
@article{zhang2016super,
  title={Super-sensitivity multiphoton frequency-domain fluorescence lifetime imaging microscopy},
  author={Zhang, Yide and Khan, Aamir A and Vigil, Genevieve D and Howard, Scott S},
  journal={Optics Express},
  volume={24},
  number={18},
  pages={20862--20867},
  year={2016},
  publisher={Optical Society of America}
}


@book{MATLAB:2019,
year = {2019},
author = {MATLAB},
title = {9.7.0.1190202 (R2019b)},
publisher = {The MathWorks Inc.},
address = {Natick, Massachusetts}
}

%vm arxiv
@article{mannam2020performance,
  title={Performance analysis of semi-supervised learning in the small-data regime using {VAE}s},
  author={Mannam, Varun and Kazemi, Arman},
  journal={arXiv preprint arXiv:2002.12164},
  year={2020}
}


%sos
@article{zhang2018super,
  title={Super-resolution fluorescence microscopy by stepwise optical saturation},
  author={Zhang, Yide and Nallathamby, Prakash D and Vigil, Genevieve D and Khan, Aamir A and Mason, Devon E and Boerckel, Joel D and Roeder, Ryan K and Howard, Scott S},
  journal={Biomedical Optics Express},
  volume={9},
  number={4},
  pages={1613--1629},
  year={2018},
  publisher={Optical Society of America}
}

%gsos
@article{zhang2018generalized,
  title={Generalized stepwise optical saturation enables super-resolution fluorescence lifetime imaging microscopy},
  author={Zhang, Yide and Benirschke, David and Abdalsalam, Ola and Howard, Scott S},
  journal={Biomedical Optics Express},
  volume={9},
  number={9},
  pages={4077--4093},
  year={2018},
  publisher={Optical Society of America}
}


%srrf
@article{gustafsson2016fast,
  title={Fast live-cell conventional fluorophore nanoscopy with {ImageJ} through super-resolution radial fluctuations},
  author={Gustafsson, Nils and Culley, Si{\^a}n and Ashdown, George and Owen, Dylan M and Pereira, Pedro Matos and Henriques, Ricardo},
  journal={Nature Communications},
  volume={7},
  number={1},
  pages={1--9},
  year={2016},
  publisher={Nature Publishing Group}
}

%yinhao basepaper
@article{zhu2018bayesian,
  title={Bayesian deep convolutional encoder--decoder networks for surrogate modeling and uncertainty quantification},
  author={Zhu, Yinhao and Zabaras, Nicholas},
  journal={Journal of Computational Physics},
  volume={366},
  pages={415--447},
  year={2018},
  publisher={Elsevier}
}


%sr paper: 
@article{DeepSTORM,
  title={Deep-{STORM}: super-resolution single-molecule microscopy by deep learning},
  author={Nehme, Elias and Weiss, Lucien E and Michaeli, Tomer and Shechtman, Yoav},
  journal={Optica},
  volume={5},
  number={4},
  pages={458--464},
  year={2018},
  publisher={Optical Society of America}
}

@inproceedings{UNet,
  title={U-{N}et: Convolutional networks for biomedical image segmentation},
  author={Ronneberger, Olaf and Fischer, Philipp and Brox, Thomas},
  booktitle={International Conference on Medical Image Computing and Computer-Assisted Intervention},
  pages={234--241},
  year={2015},
  organization={Springer}
}


%literature
@article{wang2019deep_gan1,
  title={Deep learning enables cross-modality super-resolution in fluorescence microscopy},
  author={Wang, Hongda and Rivenson, Yair and Jin, Yiyin and Wei, Zhensong and Gao, Ronald and G{\"u}nayd{\i}n, Harun and Bentolila, Laurent A and Kural, Comert and Ozcan, Aydogan},
  journal={Nature Methods},
  volume={16},
  number={1},
  pages={103--110},
  year={2019},
  publisher={Nature Publishing Group}
}

@article{zhang2019high_gan2,
  title={High-throughput, high-resolution deep learning microscopy based on registration-free generative adversarial network},
  author={Zhang, Hao and Fang, Chunyu and Xie, Xinlin and Yang, Yicong and Mei, Wei and Jin, Di and Fei, Peng},
  journal={Biomedical Optics Express},
  volume={10},
  number={3},
  pages={1044--1063},
  year={2019},
  publisher={Optical Society of America}
}

@article{ouyang2018deep_gan3,
  title={Deep learning massively accelerates super-resolution localization microscopy},
  author={Ouyang, Wei and Aristov, Andrey and Lelek, Micka{\"e}l and Hao, Xian and Zimmer, Christophe},
  journal={Nature {B}iotechnology},
  volume={36},
  number={5},
  pages={460--468},
  year={2018},
  publisher={Nature Publishing Group}
}

@article{ayas2020microscopic_fc2,
  title={Microscopic image super resolution using deep convolutional neural networks},
  author={Ayas, Selen and Ekinci, Murat},
  journal={Multimedia Tools and Applications},
  volume={79},
  number={21},
  pages={15397--15415},
  year={2020},
  publisher={Springer}
}

@article{xie2018microscopy_fc3,
  title={Microscopy cell counting and detection with fully convolutional regression networks},
  author={Xie, Weidi and Noble, J Alison and Zisserman, Andrew},
  journal={Computer Methods in Biomechanics and Biomedical Engineering: Imaging \& Visualization},
  volume={6},
  number={3},
  pages={283--292},
  year={2018},
  publisher={Taylor \& Francis}
}

@article{huszka2019super,
  title={Super-resolution optical imaging: A comparison},
  author={Huszka, Gergely and Gijs, Martin AM},
  journal={Micro and Nano Engineering},
  volume={2},
  pages={7--28},
  year={2019},
  publisher={Elsevier}
}


@article{SRRF_Nature,
  title={Fast live-cell conventional fluorophore nanoscopy with {I}mage{J} through super-resolution radial fluctuations},
  author={Gustafsson, Nils and Culley, Si{\^a}n and Ashdown, George and Owen, Dylan M and Pereira, Pedro Matos and Henriques, Ricardo},
  journal={Nature Communications},
  volume={7},
  pages={12471},
  year={2016}
}


%ml papers
@inproceedings{long2015fully,
  title={Fully convolutional networks for semantic segmentation},
  author={Long, Jonathan and Shelhamer, Evan and Darrell, Trevor},
  booktitle={Proceedings of the IEEE Conference on Computer Vision and Pattern Recognition (CVPR)},
  pages={3431--3440},
  year={2015}
}

@inproceedings{huang2017densely,
  title={Densely connected convolutional networks},
  author={Huang, Gao and Liu, Zhuang and Van Der Maaten, Laurens and Weinberger, Kilian Q},
  booktitle={Proceedings of the IEEE Conference on Computer Vision and Pattern Recognition (CVPR)},
  pages={4700--4708},
  year={2017}
}

%gan intro
@inproceedings{osokin2017gans,
  title={{GANs} for biological image synthesis},
  author={Osokin, Anton and Chessel, Anatole and Carazo Salas, Rafael E and Vaggi, Federico},
  booktitle={Proceedings of the IEEE International Conference on Computer Vision (ICCV)},
  pages={2233--2242},
  year={2017}
}

%yide new version v2
@article{instantFLIM,
  title={High-speed, long-term, {4D} \textit{in vivo} lifetime imaging in intact and injured zebrafish and mouse brains by instant {FLIM}},
  author={Zhang, Yide and Guldner, Ian H and Nichols, Evan L and Benirschke, David and Smith, Cody J and Zhang, Siyuan and Howard, Scott S},
  journal={bioRxiv},
  year={2020},
  publisher={Cold Spring Harbor Laboratory}
}

%denseED inspiration
@inproceedings{jegou2017one,
  title={The {O}ne {H}undred {L}ayers {T}iramisu: {F}ully {C}onvolutional {D}ensenets for {S}emantic {S}egmentation},
  author={J{\'e}gou, Simon and Drozdzal, Michal and Vazquez, David and Romero, Adriana and Bengio, Yoshua},
  booktitle={Proceedings of the IEEE Conference on Computer Vision and Pattern Recognition Workshops (CVPR)},
  pages={11--19},
  year={2017}
}

%ct xray
@article{kang2017deep,
  title={A deep convolutional neural network using directional wavelets for low-dose {X-ray} {CT} reconstruction},
  author={Kang, Eunhee and Min, Junhong and Ye, Jong Chul},
  journal={Medical Physics},
  volume={44},
  number={10},
  pages={e360--e375},
  year={2017},
  publisher={Wiley Online Library}
}

%ssim
@article{wang2004image,
  title={Image quality assessment: from error visibility to structural similarity},
  author={Wang, Zhou and Bovik, Alan C and Sheikh, Hamid R and Simoncelli, Eero P},
  journal={IEEE Transactions on Image Processing (TIP)},
  volume={13},
  number={4},
  pages={600--612},
  year={2004},
  publisher={IEEE}
}

%rsp
@article{laine2019nanoj,
  title={Nano{J}: a high-performance open-source super-resolution microscopy toolbox},
  author={Laine, Romain F and Tosheva, Kalina L and Gustafsson, Nils and Gray, Robert DM and Almada, Pedro and Albrecht, David and Risa, Gabriel T and Hurtig, Fredrik and Lind{\aa}s, Ann-Christin and Baum, Buzz and others},
  journal={Journal of Physics D: Applied Physics},
  volume={52},
  number={16},
  pages={163001},
  year={2019},
  publisher={IOP Publishing}
}

%sr metrics
@inproceedings{lukevs2013performance,
  title={Performance evaluation of image quality metrics with respect to their use for super-resolution enhancement},
  author={Luke{\v{s}}, Tom{\'a}{\v{s}} and Fliegel, Karel and Kl{\'\i}ma, Milo{\v{s}}},
  booktitle={2013 Fifth International Workshop on Quality of Multimedia Experience (QoMEX)},
  pages={42--43},
  year={2013},
  organization={IEEE}
}

%vm_cnn_flim
@inproceedings{mannam2021convolutional,
  title={Convolutional neural network denoising in fluorescence lifetime imaging microscopy ({FLIM})},
  author={Mannam, Varun and Zhang, Yide and Yuan, Xiaotong and Hato, Takashi and Dagher, Pierre C and Nichols, Evan L and Smith, Cody J and Dunn, Kenneth W and Howard, Scott},
  booktitle={Multiphoton Microscopy in the Biomedical Sciences XXI},
  volume={11648},
  pages={116481C},
  year={2021},
  organization={International Society for Optics and Photonics (SPIE Photonics West)}
}


%thunderstorm
@article{ovesny2014thunderstorm,
  title={Thunder{STORM}: a comprehensive {ImageJ} plug-in for {PALM} and {STORM} data analysis and super-resolution imaging},
  author={Ovesn{\`y}, Martin and K{\v{r}}{\'\i}{\v{z}}ek, Pavel and Borkovec, Josef and {\v{S}}vindrych, Zden{\v{e}}k and Hagen, Guy M},
  journal={Bioinformatics},
  volume={30},
  number={16},
  pages={2389--2390},
  year={2014},
  publisher={Oxford University Press}
}


%SAX microscopy
@article{humpolivckova2009optical,
  title={Optical saturation as a versatile tool to enhance resolution in confocal microscopy},
  author={Humpol{\'\i}{\v{c}}kov{\'a}, Jana and Benda, Ale{\v{s}} and Enderlein, J{\"o}rg},
  journal={Biophysical Journal},
  volume={97},
  number={9},
  pages={2623--2629},
  year={2009},
  publisher={Elsevier}
}

@article{fujita2007high,
  title={High-resolution confocal microscopy by saturated excitation of fluorescence},
  author={Fujita, Katsumasa and Kobayashi, Minoru and Kawano, Shogo and Yamanaka, Masahito and Kawata, Satoshi},
  journal={Physical Review Letters (PRL)},
  volume={99},
  number={22},
  pages={228105},
  year={2007},
  publisher={APS}
}

@article{cianci2004saturation,
  title={Saturation modified point spread functions in two-photon microscopy},
  author={Cianci, Gianguido C and Wu, Jianrong and Berland, Keith M},
  journal={Microscopy Research and Technique},
  volume={64},
  number={2},
  pages={135--141},
  year={2004},
  publisher={Wiley Online Library}
}


@article{weigert2018content,
  title={Content-aware image restoration: pushing the limits of fluorescence microscopy},
  author={Weigert, Martin and Schmidt, Uwe and Boothe, Tobias and M{\"u}ller, Andreas and Dibrov, Alexandr and Jain, Akanksha and Wilhelm, Benjamin and Schmidt, Deborah and Broaddus, Coleman and Culley, Si{\^a}n and others},
  journal={Nature Methods},
  volume={15},
  number={12},
  pages={1090--1097},
  year={2018},
  publisher={Nature Publishing Group}
}

%n2void
@inproceedings{krull2019noise2void,
  title={Noise2Void-learning denoising from single noisy images},
  author={Krull, Alexander and Buchholz, Tim-Oliver and Jug, Florian},
  booktitle={Proceedings of the IEEE Conference on Computer Vision and Pattern Recognition (CVPR)},
  pages={2129--2137},
  year={2019}
}


@book{webb2017introduction,
  title={Introduction to {B}iomedical {I}maging},
  author={Webb, Andrew G},
  year={2017},
  publisher={John Wiley \& Sons}
}
@article{weissleder2015advancing,
  title={Advancing biomedical imaging},
  author={Weissleder, Ralph and Nahrendorf, Matthias},
  journal={Proceedings of the National Academy of Sciences (PNAS)},
  volume={112},
  number={47},
  pages={14424--14428},
  year={2015},
  publisher={National Acad Sciences}
}

@inproceedings{liang2011optical,
  title={Optical design for biomedical imaging},
  author={Liang, Rongguang},
  year={2011},
  organization={SPIE}
}

@book{bronzino2000biomedical,
  title={Biomedical Engineering Handbook},
  author={Bronzino, Joseph D},
  volume={2},
  year={2000},
  publisher={Springer Science \& Business Media}
}

%TBME papers
@article{nie2018medical,
  title={Medical image synthesis with deep convolutional adversarial networks},
  author={Nie, Dong and Trullo, Roger and Lian, Jun and Wang, Li and Petitjean, Caroline and Ruan, Su and Wang, Qian and Shen, Dinggang},
  journal={IEEE Transactions on Biomedical Engineering (TBME)},
  volume={65},
  number={12},
  pages={2720--2730},
  year={2018},
  publisher={IEEE}
}


@article{kiranyaz2015real,
  title={Real-time patient-specific {ECG} classification by 1-{D} convolutional neural networks},
  author={Kiranyaz, Serkan and Ince, Turker and Gabbouj, Moncef},
  journal={IEEE Transactions on Biomedical Engineering (TBME)},
  volume={63},
  number={3},
  pages={664--675},
  year={2015},
  publisher={IEEE}
}

%MRI super-resol low data
@article{chun2019mri,
  title={{MRI} super-resolution reconstruction for {MRI}-guided adaptive radiotherapy using cascaded deep learning: In the presence of limited training data and unknown translation model},
  author={Chun, Jaehee and Zhang, Hao and Gach, H Michael and Olberg, Sven and Mazur, Thomas and Green, Olga and Kim, Taeho and Kim, Hyun and Kim, Jin Sung and Mutic, Sasa and others},
  journal={Medical Physics},
  volume={46},
  number={9},
  pages={4148--4164},
  year={2019},
  publisher={Wiley Online Library}
}

@article{robinson2010new,
  title={New applications of super-resolution in medical imaging},
  author={Robinson, M Dirk and Chiu, Stephanie J and Lo, J and Toth, C and Izatt, J and Farsiu, Sina},
  journal={Super-Resolution Imaging},
  volume={2010},
  pages={384--412},
  year={2010},
  publisher={CRC Press}
}
@article{yang2018low,
  title={Low-dose {X}-ray tomography through a deep convolutional neural network},
  author={Yang, Xiaogang and De Andrade, Vincent and Scullin, William and Dyer, Eva L and Kasthuri, Narayanan and De Carlo, Francesco and G{\"u}rsoy, Do{\u{g}}a},
  journal={Scientific Reports},
  volume={8},
  number={1},
  pages={1--13},
  year={2018},
  publisher={Nature Publishing Group}
}

@article{umehara2018application,
  title={Application of super-resolution convolutional neural network for enhancing image resolution in chest {CT}},
  author={Umehara, Kensuke and Ota, Junko and Ishida, Takayuki},
  journal={Journal of Digital Imaging},
  volume={31},
  number={4},
  pages={441--450},
  year={2018},
  publisher={Springer}
}


%instant_flim_new
@article{instantFLIM_new,
  title={Instant {FLIM} enables 4{D} \textit{in vivo} lifetime imaging of intact and injured zebrafish and mouse brains},
  author={Yide Zhang and Ian Guldner and Evan Nichols and David Benirschke and Cody Smith and Siyuan Zhang and Scott Howard},
  journal={Optica},
  volume={8},
  number={},
  pages={},
  year={2021},
  url = {https://www.osapublishing.org/optica/abstract.cfm?doi=10.1364/OPTICA.426870},
  doi = {10.1364/OPTICA.426870},
  publisher={Optical Society of America}
}

%w2s dataset
@inproceedings{zhou2020w2s,
  title={{W2S}: microscopy data with joint denoising and super-resolution for widefield to {SIM} mapping},
  author={Zhou, Ruofan and El Helou, Majed and Sage, Daniel and Laroche, Thierry and Seitz, Arne and S{\"u}sstrunk, Sabine},
  booktitle={European Conference on Computer Vision (ECCV)},
  pages={474--491},
  year={2020},
  organization={Springer}
}

%vm: 270622
@article{mannam2022real,
  title={Real-time image denoising of mixed {P}oisson--{G}aussian noise in fluorescence microscopy images using {ImageJ}},
  author={Mannam, Varun and Zhang, Yide and Zhu, Yinhao and Nichols, Evan and Wang, Qingfei and Sundaresan, Vignesh and Zhang, Siyuan and Smith, Cody and Bohn, Paul W and Howard, Scott S},
  journal={Optica},
  volume={9},
  number={4},
  pages={335--345},
  year={2022},
  publisher={Optica Publishing Group}
}

@article{abbe1873posts,
  title={Contributions to the theory of the microscope and microscopic perception},
  author={Abbe, Ernst},
  journal={Archive for Microscopic Anatomy},
  volume={9},
  number={1},
  pages={413--468},
  year={1873},
  publisher={Knight}
}

%adam
@article{kingma2014adam,
  title={Adam: A method for stochastic optimization},
  author={Kingma, Diederik P and Ba, Jimmy},
  journal={arXiv:1412.6980},
  year={2014}
}

%ssim vs psnr
@inproceedings{hore2010image,
  title={Image quality metrics: {PSNR} vs. {SSIM}},
  author={Hore, Alain and Ziou, Djemel},
  booktitle={2010 20th International Conference on Pattern Recognition},
  pages={2366--2369},
  year={2010},
  organization={IEEE}
}

%literature
@article{li2018dlbi,
  title={DLBI: deep learning guided Bayesian inference for structure reconstruction of super-resolution fluorescence microscopy},
  author={Li, Yu and Xu, Fan and Zhang, Fa and Xu, Pingyong and Zhang, Mingshu and Fan, Ming and Li, Lihua and Gao, Xin and Han, Renmin},
  journal={Bioinformatics},
  volume={34},
  number={13},
  pages={i284--i294},
  year={2018},
  publisher={Oxford University Press}
}

@article{zhao2022sparse,
  title={Sparse deconvolution improves the resolution of live-cell super-resolution fluorescence microscopy},
  author={Zhao, Weisong and Zhao, Shiqun and Li, Liuju and Huang, Xiaoshuai and Xing, Shijia and Zhang, Yulin and Qiu, Guohua and Han, Zhenqian and Shang, Yingxu and Sun, De-en and others},
  journal={Nature Biotechnology},
  volume={40},
  number={4},
  pages={606--617},
  year={2022},
  publisher={Nature Publishing Group}
}

@article{kaderuppan2022net,
  title={{O-Net}: A Fast and Precise Deep-Learning Architecture for Computational Super-Resolved Phase-Modulated Optical Microscopy},
  author={Kaderuppan, Shiraz S and Wong, Wai Leong Eugene and Sharma, Anurag and Woo, Wai Lok},
  journal={Microscopy and Microanalysis},
  pages={1--15},
  year={2022},
  publisher={Cambridge University Press}
}

@article{zelger2018three,
  title={Three-dimensional localization microscopy using deep learning},
  author={Zelger, P and Kaser, K and Rossboth, B and Velas, L and Sch{\"u}tz, GJ and Jesacher, A},
  journal={Optics express},
  volume={26},
  number={25},
  pages={33166--33179},
  year={2018},
  publisher={Optical Society of America}
}


%vm
@article{hemdan2020covidx,
  title={Covid{X}-net: A framework of deep learning classifiers to diagnose {Covid-19} in {X-ray} images},
  author={Hemdan, Ezz El-Din and Shouman, Marwa A and Karar, Mohamed Esmail},
  journal={arXiv preprint arXiv:2003.11055},
  year={2020}
}
@article{jain2021deep,
  title={Deep learning based detection and analysis of {COVID-19} on chest {X-ray} images},
  author={Jain, Rachna and Gupta, Meenu and Taneja, Soham and Hemanth, D Jude},
  journal={Applied Intelligence},
  volume={51},
  number={3},
  pages={1690--1700},
  year={2021},
  publisher={Springer}
}
@article{baltruschat2019comparison,
  title={Comparison of deep learning approaches for multi-label chest {X-ray} classification},
  author={Baltruschat, Ivo M and Nickisch, Hannes and Grass, Michael and Knopp, Tobias and Saalbach, Axel},
  journal={Scientific Reports},
  volume={9},
  number={1},
  pages={1--10},
  year={2019},
  publisher={Nature Publishing Group}
}
@article{liu2018applications,
  title={Applications of deep learning to {MRI} images: A survey},
  author={Liu, Jin and Pan, Yi and Li, Min and Chen, Ziyue and Tang, Lu and Lu, Chengqian and Wang, Jianxin},
  journal={Big Data Mining and Analytics},
  volume={1},
  number={1},
  pages={1--18},
  year={2018},
  publisher={TUP}
}


%new papers for reference
@inproceedings{shocher2018zero,
  title={“zero-shot” super-resolution using deep internal learning},
  author={Shocher, Assaf and Cohen, Nadav and Irani, Michal},
  booktitle={Proceedings of the IEEE Conference on Computer Vision and Pattern Recognition (CVPR)},
  pages={3118--3126},
  year={2018}
}
@inproceedings{soh2020meta,
  title={Meta-transfer learning for zero-shot super-resolution},
  author={Soh, Jae Woong and Cho, Sunwoo and Cho, Nam Ik},
  booktitle={Proceedings of the IEEE/CVF Conference on Computer Vision and Pattern Recognition (CVPR)},
  pages={3516--3525},
  year={2020}
}
@inproceedings{cheng2021light,
  title={Light field super-resolution with zero-shot learning},
  author={Cheng, Zhen and Xiong, Zhiwei and Chen, Chang and Liu, Dong and Zha, Zheng-Jun},
  booktitle={Proceedings of the IEEE/CVF Conference on Computer Vision and Pattern Recognition (CVPR)},
  pages={10010--10019},
  year={2021}
}
@article{set5dataset,
  title={Low-complexity single-image super-resolution based on nonnegative neighbor embedding},
  author={Bevilacqua, Marco and Roumy, Aline and Guillemot, Christine and Alberi-Morel, Marie Line},
  year={2012},
  publisher={{BMVA} Press}
}
@inproceedings{bsd100dataset,
  title={A database of human segmented natural images and its application to evaluating segmentation algorithms and measuring ecological statistics},
  author={Martin, David and Fowlkes, Charless and Tal, Doron and Malik, Jitendra},
  booktitle={Proceedings Eighth {IEEE} International Conference on Computer Vision. {ICCV} 2001},
  volume={2},
  pages={416--423},
  year={2001},
  organization={IEEE}
}
@inproceedings{div2kdataset,
  title={Ntire 2017 challenge on single image super-resolution: Dataset and study},
  author={Agustsson, Eirikur and Timofte, Radu},
  booktitle={Proceedings of the {IEEE} Conference on Computer Vision and Pattern Recognition Workshops (CVPR)},
  pages={126--135},
  year={2017}
}
@article{wang2020deep,
  title={Deep learning for image super-resolution: A survey},
  author={Wang, Zhihao and Chen, Jian and Hoi, Steven CH},
  journal={IEEE Transactions on Pattern Analysis and Machine Intelligence},
  volume={43},
  number={10},
  pages={3365--3387},
  year={2020},
  publisher={IEEE}
}
@article{cheng2021one,
  title={“One-Shot” Super-Resolution via Backward Style Transfer for Fast High-Resolution Style Transfer},
  author={Cheng, Jikang and Han, Zhen and Wang, Zhongyuan and Chen, Liang},
  journal={IEEE Signal Processing Letters},
  volume={28},
  pages={1485--1489},
  year={2021},
  publisher={IEEE}
}
@article{tang2021lightweight,
  title={Lightweight network with one-shot aggregation for image super-resolution},
  author={Tang, Rui and Chen, Lihui and Zou, Yiye and Lai, Zhibing and Albertini, Marcelo Keese and Yang, Xiaomin},
  journal={Journal of Real-Time Image Processing},
  volume={18},
  number={4},
  pages={1275--1284},
  year={2021},
  publisher={Springer}
}
@article{wei2020boosting,
  title={Boosting one-shot spectral super-resolution using transfer learning},
  author={Wei, Wei and Sun, Yuxuan and Zhang, Lei and Nie, Jiangtao and Zhang, Yanning},
  journal={IEEE Transactions on Computational Imaging},
  volume={6},
  pages={1459--1470},
  year={2020},
  publisher={IEEE}
}

@misc{mannam2022overcoming,
  title={Overcoming Fundamental Limits of Three-Dimensional \textit{{in vivo}} Fluorescence Imaging Using Machine Learning},
  author={Mannam, Varun},
  year={2022},
  url={https://curate.nd.edu/show/5x21td99n58}, 
  DOI={10.7274/5x21td99n58}, 
  publisher={University of Notre Dame}
}

@inproceedings{wang2018esrgan,
  title={{ESRGAN}: Enhanced super-resolution generative adversarial networks},
  author={Wang, Xintao and Yu, Ke and Wu, Shixiang and Gu, Jinjin and Liu, Yihao and Dong, Chao and Qiao, Yu and Change Loy, Chen},
  booktitle={Proceedings of the European Conference on Computer Vision Workshops (ECCV)},
  pages={0--0},
  year={2018}
}
@article{nasrollahi2020deep,
  title={Deep artifact-free residual network for single-image super-resolution},
  author={Nasrollahi, Hamdollah and Farajzadeh, Kamran and Hosseini, Vahid and Zarezadeh, Esmaeil and Abdollahzadeh, Milad},
  journal={Signal, Image and Video Processing},
  volume={14},
  number={2},
  pages={407--415},
  year={2020},
  publisher={Springer}
}
@book{yeh2019computational,
  title={Computational fluorescence and phase super-resolution microscopy},
  author={Yeh, Li-Hao},
  year={2019},
  publisher={eScholarship, University of California}
}

@article{fish1995blind,
  title={Blind deconvolution by means of the Richardson--Lucy algorithm},
  author={Fish, DA and Brinicombe, AM and Pike, ER and Walker, JG},
  journal={JOSA A},
  volume={12},
  number={1},
  pages={58--65},
  year={1995},
  publisher={Optica Publishing Group}
}
@inproceedings{mannam2022deconvolution,
  title={Deconvolution of fluorescence lifetime imaging microscopy {(FLIM)}},
  author={Mannam, Varun and Yuan, Xiaotong and Howard, Scott},
  booktitle={Multiphoton Microscopy in the Biomedical Sciences XXII},
  volume={11965},
  pages={38--43},
  year={2022},
  organization={SPIE}
}
@inproceedings{baeza2017quality,
  title={Quality-efficiency trade-offs in machine learning for text processing},
  author={Baeza-Yates, Ricardo and Liaghat, Zeinab},
  booktitle={2017 IEEE International Conference on BigData ({BigData})},
  pages={897--904},
  year={2017},
  organization={IEEE}
}
@inproceedings{feng2021gans,
  title={When do GANs replicate? On the choice of dataset size},
  author={Feng, Qianli and Guo, Chenqi and Benitez-Quiroz, Fabian and Martinez, Aleix M},
  booktitle={Proceedings of the IEEE/CVF International Conference on Computer Vision (ICCV)},
  pages={6701--6710},
  year={2021}
}
@inproceedings{linjordet2019impact,
  title={Impact of training dataset size on neural answer selection models},
  author={Linjordet, Trond and Balog, Krisztian},
  booktitle={European Conference on Information Retrieval},
  pages={828--835},
  year={2019},
  organization={Springer}
}
@article{hosseinzadeh2022deep,
  title={Deep learning--assisted prostate cancer detection on bi-parametric MRI: minimum training data size requirements and effect of prior knowledge},
  author={Hosseinzadeh, Matin and Saha, Anindo and Brand, Patrick and Slootweg, Ilse and de Rooij, Maarten and Huisman, Henkjan},
  journal={European Radiology},
  volume={32},
  number={4},
  pages={2224--2234},
  year={2022},
  publisher={Springer}
}

@misc{trainingdatasetsize,
  title={Impact of Dataset Size on Deep Learning Model Skill And Performance Estimates},
  author={Brownlee, Jason},
  year={2019},
  url={https://machinelearningmastery.com/impact-of-dataset-size-on-deep-learning-model-skill-and-performance-estimates/}, 
publisher={online}
}
@article{luo2020super,
  title={Super-resolution structured illumination microscopy reconstruction using a least-squares solver},
  author={Luo, Jintao and Li, Chuankang and Liu, Qiulan and Wu, Junling and Li, Haifeng and Kuang, Cuifang and Hao, Xiang and Liu, Xu},
  journal={Frontiers in Physics},
  pages={118},
  year={2020},
  publisher={Frontiers}
}

@article{prakash2022super,
  title={Super-resolution microscopy: a brief history and new avenues},
  author={Prakash, Kirti and Diederich, Benedict and Heintzmann, Rainer and Schermelleh, Lothar},
  journal={Philosophical Transactions of the Royal Society A},
  volume={380},
  number={2220},
  pages={20210110},
  year={2022},
  publisher={The Royal Society}
}

@inproceedings{glorot2011deep,
  title={Deep sparse rectifier neural networks},
  author={Glorot, Xavier and Bordes, Antoine and Bengio, Yoshua},
  booktitle={Proceedings of the fourteenth international conference on artificial intelligence and statistics},
  pages={315--323},
  year={2011},
  organization={JMLR Workshop and Conference Proceedings}
}
@inproceedings{ioffe2015batch,
  title={Batch normalization: Accelerating deep network training by reducing internal covariate shift},
  author={Ioffe, Sergey and Szegedy, Christian},
  booktitle={International Conference on Machine Learning (ICML)},
  pages={448--456},
  year={2015},
  organization={PMLR}
}

%gans: 
@inproceedings{chen2018image,
  title={Image blind denoising with generative adversarial network based noise modeling},
  author={Chen, Jingwen and Chen, Jiawei and Chao, Hongyang and Yang, Ming},
  booktitle={Proceedings of the IEEE Conference on Computer Vision and Pattern Recognition (CVPR)},
  pages={3155--3164},
  year={2018}
}
@article{agarwal2016multiple,
  title={Multiple signal classification algorithm for super-resolution fluorescence microscopy},
  author={Agarwal, Krishna and Mach{\'a}{\v{n}}, Radek},
  journal={Nature Communications},
  volume={7},
  number={1},
  pages={1--9},
  year={2016},
  publisher={Nature Publishing Group}
}
@article{park2022deep,
  title={Deep learning enables reference-free isotropic super-resolution for volumetric fluorescence microscopy},
  author={Park, Hyoungjun and Na, Myeongsu and Kim, Bumju and Park, Soohyun and Kim, Ki Hean and Chang, Sunghoe and Ye, Jong Chul},
  journal={Nature Communications},
  volume={13},
  number={1},
  pages={1--12},
  year={2022},
  publisher={Nature Publishing Group}
}
@article{liao2022deep,
  title={Deep-learning-based methods for super-resolution fluorescence microscopy.},
  author={Liao, Jianhui and Qu, Junle and Hao, Yongqi and Li, Jia},
  journal={Journal of Innovative Optical Health Sciences},
  year={2022}
}
@article{huang2023enhancing,
  title={Enhancing image resolution of confocal fluorescence microscopy with deep learning},
  author={Huang, Boyi and Li, Jia and Yao, Bowen and Yang, Zhigang and Lam, Edmund Y and Zhang, Jia and Yan, Wei and Qu, Junle},
  journal={PhotoniX},
  volume={4},
  number={1},
  pages={1--22},
  year={2023},
  publisher={SpringerOpen}
}
@misc{waters2009accuracy,
  title={Accuracy and precision in quantitative fluorescence microscopy},
  author={Waters, Jennifer C},
  year={2009},
  publisher={The Rockefeller University Press}
}
@inproceedings{mannam2022low,
  title={Low dosage {3D} volume fluorescence microscopy imaging using compressive sensing},
  author={Mannam, Varun and Brandt, Jacob and Smith, Cody J and Howard, Scott},
  booktitle={Three-Dimensional and Multidimensional Microscopy: Image Acquisition and Processing XXIX},
  volume={11966},
  pages={7--13},
  year={2022},
  organization={SPIE}
}
@article{howard2020packet,
  title={Packet compressed sensing imaging ({PCSI}): Robust image transmission over noisy channels},
  author={Howard, Scott and Barthelmes, Grant and Ravasio, Cara and Huang, Lisa and Poag, Benjamin and Mannam, Varun},
  journal={arXiv preprint arXiv:2009.11455},
  year={2020}
}
