## Supplementary material for "Small Training Dataset Convolutional Neural Networks for Application Specific Super-Resolution Microscopy": super-resolution latex file: SRDenseED_manuscript.tex

\documentclass[12pt]{spieman}  %>>> use for US letter paper
%\documentclass[a4paper]{spie}  %>>> use this instead for A4 paper
%\documentclass[nocompress]{spie}  %>>> to avoid compression of citations
\usepackage{setspace}
\usepackage{tocloft}
%\usepackage{footnotebackref}
\renewcommand{\baselinestretch}{1.0} % Change to 1.65 for double spacing
\usepackage{amsmath,amsfonts,amssymb}
\usepackage{graphicx}
\usepackage[colorlinks=true, allcolors=blue]{hyperref}
\usepackage{multirow}
\usepackage{upgreek}
\usepackage{gensymb}
\usepackage{mathtools} %for multi-lines 
\usepackage[english]{babel}
\usepackage{lastpage}
\pagestyle{plain}
\pagenumbering{arabic}
% \usepackage{footnotebackref}
\usepackage{hyperref}
\usepackage{rotating} %rotate table
\usepackage{caption, tabularx}
\usepackage{enumitem, ragged2e}
\usepackage[svgnames]{xcolor}
\usepackage{csquotes}
\usepackage{appendix}
\usepackage{siunitx}

\usepackage{xurl} %long url break into parts
%\usepackage[demo]{graphicx}
%\usepackage{caption}
\usepackage{subcaption}
\usepackage[normalem]{ulem}
\newcommand{\minitab}[2][l]{\begin{tabular}{#1}#2\end{tabular}}
%\title{Ultra-small training dataset neural networks for biomedical imaging} %change later old one
\title{Small Training Dataset Convolutional Neural Networks for Application Specific Super-Resolution Microscopy}
\author[1]{Varun Mannam$^*$}
\author[1]{Scott Howard$^*$}
\affil[1]{Department of Electrical Engineering, University of Notre Dame, Notre Dame, IN 46556, USA}

% Option to view page numbers
\pagestyle{plain} % change to \pagestyle{plain} for page numbers   
\setcounter{page}{1} % Set start page numbering at e.g. 301
%page numbers
\usepackage{lineno}
\linenumbers
\renewcommand{\cftdotsep}{\cftnodots}
\cftpagenumbersoff{figure}
\cftpagenumbersoff{table} 
\newcommand{\cc}[1]{\textcolor{black}{#1}}%blue


\begin{document} 
\maketitle
%\dates{Compiled \today}

\begin{abstract} \newline
\textbf{Significance:}
Machine learning (ML) models based on deep convolutional neural networks have been used to significantly increase microscopy resolution, speed (signal-to-noise ratio), and data interpretation. The bottleneck in developing effective ML systems is often the need to acquire large datasets to train the neural network. This paper demonstrates how adding a ``dense encoder-decoder'' block can be used to effectively train a neural network  that produces super-resolution images from conventional microscopy diffraction-limited images trained using a small dataset (15 field-of-views).
\newline
\textbf{Aim:} ML helps to retrieve super-resolution information from a diffraction-limited image when trained with a massive training dataset. The aim of this work is to demonstrate a neural network that estimates super-resolution images from diffraction-limited images using  modifications that enable training with a small dataset.
\newline
\textbf{Approach:} We employ ``Dense Encoder-Decoder" (called DenseED) blocks in existing super-resolution ML network architectures. DenseED blocks use a dense layer that concatenates features from the previous convolutional layer to the next convolutional layer. DenseED blocks in fully convolutional networks (FCNs) estimate the super-resolution images when trained with a small training dataset (15 field-of-views) of human cells from the Widefield2SIM dataset and in fluorescent-labeled fixed bovine pulmonary artery endothelial cells (BPAE samples).
\newline
\textbf{Results:} Conventional ML models without DenseED blocks trained on small datasets fail to accurately estimate super-resolution images while models including the DenseED blocks can. The average peak signal-to-noise ratio (PSNR) and resolution improvements achieved by networks containing DenseED blocks are $\approx$3.2 dB and 2$\times$, respectively. We evaluated various configurations of target image generation methods (e.g, experimentally captured a target and computationally generated target) that are used to train FCNs with and without DenseED blocks and showed including DenseED blocks in simple FCNs outperforms compared to simple FCNs without DenseED blocks. 
\newline
\textbf{Conclusions:} DenseED blocks in neural networks show accurate extraction of super-resolution images even if the ML model is trained with a small training dataset of 15 field-of-views. This approach shows that microscopy applications can use DenseED blocks to train on smaller datasets that are application-specific imaging platforms and there is a promise for applying this to other imaging modalities such as MRI/X-ray, etc.
\newline
\end{abstract}

% Include a list of keywords after the abstract 
\keywords{small datasets, biomedical imaging, fluorescence microscopy, super-resolution, diffraction-limited, machine learning, fully convolutional networks, convolutional neural networks, generative adversarial networks, Dense Encoder-Decoder, dense layer}

% Include email contact information for the corresponding author

%%%%%%%%%%%%%%%%%%%%%%%%%%%%%%%%%%%%%%%%%%%%%%%%%%%%%%%%%%%%%%%%%%%%%%%%%%%%. 
%%%%%%%%%%%%%%%%%%%%%%%%%%%%%%% Introduction %%%%%%%%%%%%%%%%%%%%%%%%%%%%%%%
%%%%%%%%%%%%%%%%%%%%%%%%%%%%%%%%%%%%%%%%%%%%%%%%%%%%%%%%%%%%%%%%%%%%%%%%%%%%
\section{Introduction}  \label{introduction}
Significant technical advances have allowed researchers to break through the fundamental limits to biomedical imaging resolution and speed, subsequently leading to significant improvements in data analysis and interpretation \cite{bronzino2000biomedical, webb2017introduction, liang2011optical}. However, many of these approaches require specialized equipment and training, limiting their applicability. For example, the diffraction limit in fluorescence microscopy has been overcome by a wide variety of super-resolution techniques \cite{STED, SIM, STORM, PALM}. To make these technical advances more widely available, machine learning (ML) approaches have been used to estimate super-resolution images obtained from those techniques while using conventional and commonly available imaging platforms \cite{nie2018medical, DeepSTORM, mannam2022real}. These ML models are powerful and easily distributable; however, they require significantly large training datasets \cite{DeepSTORM, mannam2022real} ($\geq$10,000 images) that are often prohibitively expensive and time-consuming to generate. This limitation is especially true for biomedical imaging such as \textit{in vivo} imaging, MRI imaging, and X-ray \cite{robinson2010new, yang2018low, umehara2018application, chun2019mri}. In addition, the imaging experimental setup for the above-mentioned applications is specific to those applications (denoted as application-specific) with variance across experimental equipment. Without large training datasets, existing ML models are less accurate and not capable of generating super-resolution images from diffraction-limited images.

In this paper, we develop, demonstrate, and evaluate using a small training dataset (much less than 1000 images) with convolutional neural network (CNN) models by incorporating new dense Encoder-decoder (``DenseED") blocks \cite{huang2017densely} that can successfully estimate fluorescence microscope images with resolution enhancements. To illustrate this method, we trained a CNN with DenseED blocks using small training datasets that increased both the resolution by a factor of 2 and the peak signal-to-noise ratio (PSNR) by 3.2 dB. Such performance is not possible using conventional CNNs without DenseED blocks. The results show how ML models can be novel for specific equipment and applications using small datasets acquired by that specific tool.

%%%%%%%%%%%%%%%%%%%%%%%%%%%%%%%%%%%%%%%%%%%%%%%%%%%%%%%%%%%%%%%%%%%%%%%%%%%%
%%%%%%%%%%%%%%%%%%%%%%%%%%%%%%% Methods %%%%%%%%%%%%%%%%%%%%%%%%%%%%%%%%%%%%
%%%%%%%%%%%%%%%%%%%%%%%%%%%%%%%%%%%%%%%%%%%%%%%%%%%%%%%%%%%%%%%%%%%%%%%%%%%%
\section{Methods and Dataset Creation} \label{methods}
\subsection{Traditional super-resolution methods} \label{s21}
Fluorescence microscopy is a key research tool throughout biology \cite{lichtman2005fluorescence}. However, the spatial resolution of an image generated by conventional fluorescence microscopy is limited to a few hundred nanometers defined by the diffraction limit of light \cite{abbe1873posts}. The limited resolution hinders further observation and investigation of objects at a sub-cellular or molecular scale, such as mitochondria, microtubules, nanopores, and proteins within cells and tissues. Many fluorescence microscopy super-resolution methods can overcome the diffraction limit and achieve better resolutions up to ten times greater than conventional microscopy techniques. Experimental methods such as stimulated emission depletion (STED)  \cite{STED}, structured illumination microscopy (SIM) \cite{SIM}, and non-linear SIM \cite{NSIM, SIM_PNAS} perform super-resolution imaging; but typically, they require dedicating imaging platforms. Exploiting the non-linearity of excitation saturation in scanning microscopy enables super-resolution microscopy in conventional microscope platforms \cite{fujita2007high,humpolivckova2009optical, GSOS}. Localization and statistical approaches, including stochastic optical reconstruction microscopy (STORM) \cite{STORM} and photoactivated localization microscopy (PALM) \cite{PALM} can also enhance the image resolution but require special fluorophores and extensive computation. Computational methods such as Super-resolution radial fluctuation (SRRF) \cite{SRRF_Nature} can be used to perform super-resolution imaging. SRRF can generate images with a resolution comparable to localization approaches without requiring complicated hardware setups and special imaging conditions. Even so, it requires numerous diffraction-limited images to be collected within a single FOV and is computationally expensive. To achieve the benefits of super-resolution techniques on conventional imaging platforms, ML approaches can be used.

\subsection{ML-based super-resolution methods in literature} \label{s22}
\cc{ML has gained attention for its fast processing speed, and wide applications, such as image classification \cite{goodfellow2016deep, mannam2020machine}, image denoising \cite{mannam2022real}, image segmentation \cite{mannam2021convolutional}, and image compression \cite{mannam2022low, howard2020packet}. The ML models achieve high performance and generalization capacity when trained with a large training dataset \cite{mannam2022real}. However, obtaining a large training dataset is often prohibitively expensive or difficult \cite{zhang2019poisson}. In addition, the variance between the same models of the experimental equipment can be large (due to each application-specific equipment calibration/setup setting being different), making generalizability difficult \cite{waters2009accuracy}. Hence, the training dataset size is often limited and application specific. Nonetheless, existing ML models show high performance when trained with a large training dataset. Hence there is a trade-off between application-specific ML model performance vs. training dataset size \cite{feng2021gans, hosseinzadeh2022deep, trainingdatasetsize, baeza2017quality, linjordet2019impact}. }
%specific to super-resolution
%removed fig1 explanation 15th aug here
\begin{figure}[!ht]
\centering
\includegraphics[page=1,width=0.9\linewidth]{SRDenseED_all_figures_list.pdf}
\caption{Block diagram of fully convolutional networks with skip connections, including the encoder and decoder blocks. Here the network indicates the autoencoder (AE) and U-Net architectures without and with skip connections \cite{mannam2020instant}, respectively. Encoder and decoder blocks consist of batch-norm, ReLU (rectified linear unit), and convolution layers. Conv(s2) and Conv$^{T}$(s2) indicate the convolution and convolution transpose layers with a stride of 2, respectively. The symbol $\oplus$ represents the concatenation layer that combines the output from the encoder layer and decoder layer in the number of channels dimension.} \label{sr_fig2}
\end{figure} 

In literature, existing ML-based super-resolution methods can be classified into two categories \cite{mannam2021deep}: fully convolutional networks (FCNs) and generative adversarial networks (GANs). FCNs containing a combination of encoder and decoder blocks \cite{mannam2020performance} as shown in Fig.~\ref{sr_fig2}. \cc{Some examples of FCN architectures are U-Net \cite{UNet}, dense nets \cite{huang2017densely}, residual nets \cite{zhang2018residual}, and autoencoders (AEs) \cite{DeepSTORM}.} FCNs architecture includes multiple encoder and decoder blocks (convolutional layers), and the output is generated by combining the output of encoder layers from different convolutional layers in encoder and decoder blocks (refer to Fig.~\ref{sr_fig2}). To pass the features generated in the encoder blocks to the corresponding decoder blocks, Skip connections are helpful (refer to Fig.~\ref{sr_fig2}). GANs architecture is based on simultaneously optimizing two networks (generator and discriminator) \cite{osokin2017gans}. Two networks compete to generate the best images similar to target images from input images. In GANs, the generator network is a simple FCN (i.e., the generator consists of encoder and decoder blocks). The discriminator network consists of convolutional layers followed by the fully connected layers that generate the probability that the generator network output (estimated SR image here) looks like the real image (similar to the target image). Since the GANs generator architecture is a simple FCN architecture (to generate super-resolution images), in this paper we show our demonstrated approach using only FCN architecture. \cc{Additional details about GANs including GANs encoder and decoder can be found in these references \cite{chen2018image, li2018dlbi, agarwal2016multiple, liao2022deep, park2022deep, huang2023enhancing}. More details about the GANs including architecture, loss function, and optimization are provided in our GitHub location.}
%It is also important to achieve high performance even if the ML models are trained with a small training dataset.  
%explanation of fig1

\cc{In addition, advanced ML models such as zero-shot super-resolution (ZSSR) \cite{cheng2021light, soh2020meta, shocher2018zero, wang2020deep} and one-shot super-resolution (OSSR) \cite{tang2021lightweight, cheng2021one, wei2020boosting} with CNNs have been demonstrated to estimate high-resolution images from low-resolution ones. In the case of the ZSSR, the ML model is trained with the test image itself (hence, no-training dataset, and it is an unsupervised ML method), and performance is limited due to no training dataset. In the case of the OSSR, an extensive training dataset is used to get the high-resolution features, and ML model weights are stored. After that, a small training dataset is used to retrain the ML model with pre-trained model weights. Hence, in the OSSR case, you need two training datasets with similar features in the application-specific imaging. However, these ML models in the literature are trained on color images with datasets such as Set5 dataset \cite{set5dataset}, BSD100 images \cite{bsd100dataset}, and DIV2k images \cite{div2kdataset} but not on application-specific for example, fluorescence microscopy datasets \cite{zhang2019poisson}. Wang. et., \cite{wang2020deep} provided a consolidated summary of super-resolution methods in deep learning. In application-specific super-resolution generation, the existing computational methods that use no training data (self-supervised learning) are computationally expensive (iterative methods like image deconvolution) and lead to poor performance. In contrast, if the training dataset is large, the existing ML-based models provide higher performance, but acquiring a large training dataset (diffraction-limited and target images) is computationally expensive. Hence, finding a balance between the training dataset size and generated super-resolution images quality is significant, and this paper contributes by showing an ML-based method to mitigate the issue by providing super-resolution images accurately even if the ML model is trained with a small training dataset with input as diffraction-limited image and target as super-resolution image, respectively. Furthermore, this ML model can be applied to other application-specific super-resolution generation with a small dataset.}  

\begin{table}[!ht]
\begin{tabular}{|l|l|l|l|}
\hline
Papers  & Architecture & Training Dataset Size   & Sample details    \\ \hline
Deep-STORM \cite{DeepSTORM} & FCNs  & 7,000  & Microtubules  \\ \hline
Residual CNN's \cite{ayas2020microscopic_fc2} & FCNs  & 16,000  & Blood samples  \\ \hline
GANs Structure\cite{wang2019deep_gan1} & GANs & 2,000 (each fluorophore) & BPAE samples and Nano beads \\ \hline
RFGANs \cite{zhang2019high_gan2} & GANs  & 1,080 (increase in FOV)  & Fibroblast in mouse brain   \\ \hline
ANNA-PALM \cite{ouyang2018deep_gan3} & GANs & 30,000   & Microtubules and Nanopores  \\ \hline
\end{tabular} 
\caption{Summary of existing ML super-resolution methods with fluorescence microscopy data.} \label{ML_compare}
\end{table}

In fluorescence microscopy, traditional FCNs have been applied to generate super-resolution images from simulated and experimental data. The trained ML model (FCNs) performance is evaluated by comparing the estimated super-resolution images with the target images acquired using super-resolution microscopes. Table.~\ref{ML_compare} shows a few examples of ML models including architecture (either FCNs or GANs) and the size of the training dataset used in literature to generate fluorescence microscopy super-resolution images. In Nehme's work \cite{DeepSTORM}, the FCN architecture consists of three encoders and three decoder blocks, respectively, and is trained with 7,000 images. In Ayas's work \cite{ayas2020microscopic_fc2}, the FCN architecture includes a 20-layer residual network with blood samples trained with 16,000 images. In Wang's work \cite{wang2019deep_gan1}, the architecture is GANs with the generator network is similar to the U-Net \cite{UNet} architecture and the discriminator network consists of fully connected layers trained with 2,000 BPAE sample images for each fluorophore. Similarly, in Zhang's \cite{zhang2019high_gan2} work the ML model is GANs architecture consists of a generator network with 16-layers residual connections, and a discriminator network consists of fully connected layers with 1,080 images of fibroblast in a mouse brain. Finally, in Ouyang's work \cite{ouyang2018deep_gan3} a GANs architecture with the generator network consists of U-Net with (8,8) encoder and decoder blocks, respectively, and the discriminator network consists of fully connected layers trained with 30,000 PALM images of microtubules. Despite the ability to obtain super-resolution images from diffraction-limited images, all of the above-mentioned ML-based super-resolution models are data-driven. These trained ML models require a large training dataset (more than 1,000 images) to generate super-resolution images in fluorescence microscopy.

\subsection{FCNs with Dense Encoder Decoder} \label{s23}
This section explains the Dense Encoder-Decoder method and how it is derived from the existing FCNs architecture to provide super-resolution images when trained with a small training dataset. FCNs \cite{long2015fully} are used for pixel-wise prediction, e.g. semantic segmentation \cite{UNet}, image denoising \cite{mannam2022real}, super-resolution\cite{mannam2021deep} and low dose computer tomography X-ray reconstruction \cite{kang2017deep}. Fig.~\ref{sr_fig2} shows the FCNs architecture with encoding and decoding blocks with skip connections. The convolutional layer contains the input image convolved with a kernel that extracts particular features from the input images (for example, edges, backgrounds, and objects with different shapes). Here the number of kernels used in the convolutional layer is called the \enquote{number of feature maps} and the output of the convolution indicates the \enquote{feature map} with its dimension as \enquote{feature map size}. Typically an encoding block contains the convolutional layer with double feature maps and half of the feature map size. Encoder block is used to extract important features, thereby reducing feature map size to half. In this way, we select only the essential features as the output of the encoder block. The decoder block works exactly opposite to the encoder block; its output reduces the number of feature maps to half and doubles the feature map size. Extracting complex features such as super-resolution images from diffraction-limited images requires more encoder and decoder blocks in the ML model.

\cc{However, the feature map reaches a minimum image dimension with more encoding blocks, and super-resolution images cannot be restored using decoder blocks alone without Skip or residual or dense connections due to vanishing gradients issue in deep learning \cite{glorot2011deep, ioffe2015batch} (please refer to Fig.~\ref{sr_fig2}). In other words, coarse features are not passed through the decoder blocks in the case of deep networks. however, this requirement is not necessary when ML models contain only a small number of encoder and decoder blocks.} This minimum image dimension of the encoder is called the \enquote{latent space}. Additionally, as the number of encoder and decoder blocks increases, the number of kernel parameters (i.e., weights of the neural network) increases exponentially, which is parameter inefficient (requiring considerable computation time) ML model. As the number of encoder and decoder blocks increases, the feature map size is reduced, and the essential features are lost. Therefore, \enquote{skip connections} are introduced between encoder and decoder blocks to pass finer features (such as mitochondria and microtubules) to the decoder blocks from encoder blocks. This modified FCN architecture is called \enquote{U-Nets} \cite{UNet, mannam2022real}, which is shown in Fig.~\ref{sr_fig2} with dashed arrows, and $\oplus$ indicates the concatenation of features from the encoder block and the output of the previous decoder block. Another ML model that belongs to the FCN architecture is the ``Residual-Net" \cite{zhang2017beyond}, which consists of residual layers (or skip connection from input to output directly) where input is passed through a couple of convolutional layers. Each convolutional layer consists of convolution, non-linear elements (such as ReLU), and normalization (batch norm) layers. The last convolutional layer output is concatenated with the input. The estimated output image from the convolutional layer is the residual between target and input images (for example, noise, the subtraction of the noise input with a clear target). 

\begin{figure}[!ht]
\centering
\includegraphics[page=2,width=1.0\linewidth]{SRDenseED_all_figures_list.pdf}
\caption{Block diagram of Fully convolutional networks with dense blocks (a). dense blocks consist of multiple dense layers (b) where each dense layer's input feature maps are concatenated progressively. The dense layer (c) consists of the batch norm, ReLU, and convolution layer with stride 1 in sequence order.} \label{sr_fig3}
\end{figure}

To allow for the FCNs with higher performance when trained with the small training dataset, the modified residual connections are helpful. These modified residual connections are originally developed for physical systems and computer vision tasks, DenseED \cite{zhu2018bayesian} is the state-of-the-art CNN architecture (modified version of residual layers) due to its backbone of dense layers, which passes the extracted features from the previous layer to all next layers in a feed-forward fashion. This paper shows how to utilize these DenseED blocks to build our super-resolution ML model that works with a small dataset. Fig.~\ref{sr_fig3}(a) shows the demonstrated ML model (DenseED in FCNs) for super-resolution using an ultra-small training dataset. Fig.~\ref{sr_fig3}(a) is similar to Fig.~\ref{sr_fig2} but with additional DenseED blocks added after the encoder and decoder blocks. Fig.~\ref{sr_fig3}(b) shows the DenseED block, which consists of multiple dense layers, which is another way of passing features from one layer to the next. Dense layers \cite{huang2017densely, jegou2017one} are used to create dense connections between all layers to improve the information (gradient) flow through the complete ML model for better parameter efficiency. Fig.~\ref{sr_fig3}(c) shows the dense layer connection for $i^{th}$ dense layer with input feature maps of $x_0$ (output of the previous layer) and passed through the dense layer with output feature maps of $x_1$; total feature maps are the concatenation of input and output feature maps [$x_0$, $x_1$]. In the dense layer, the convolution operation is performed with a stride of 1. Fig.~\ref{sr_fig3}(b) shows a dense block with three dense layers, where each layer provides two feature maps as output. The dense layer establishes connections from the previous convolutional layer to all subsequent convolutional layers in the dense block. In other words, one layer's input features are concatenated to this layer's output features, which serve as the input features to the next layer. If the input has $K_0$ feature maps, and each layer of the outputs has $K$ feature maps, then the $i^{th}$ layer would have an input with $K_0 + (i*K)$ feature maps, i.e., the number of feature maps in dense block grows linearly with the depth, and $K$ here is referred to as the growth rate. More dense layers are required for the given feature map size within a dense block to access the complex features. With more dense layers in a dense block, the total output feature maps increase linearly with the growth rate $K$.

For image enhancements in FCNs, encoding and decoding blocks are required to change feature maps' size, making the concatenation of feature maps unfeasible across different feature map size blocks. Hence particular encoding and decoding blocks are used to solve this problem. A dense block contains multiple dense layers whose input and output feature maps are the same size. Each dense block has two design parameters: the number of layers $L$ and the growth rate $K$ for each layer. We consider the growth rate $K$ a constant value for all the dense blocks in our work. Here the encoding block typically is half the feature map size, while the decoding block doubles the feature map size. Both two blocks reduce the number of feature maps to half. Fig.~\ref{sr_fig3}(a) shows the complete FCNs with the DenseED (SRDenseED) ML model used to generate the super-resolution images using a small training dataset. Dense blocks, encoding blocks, and decoding blocks are marked with different colors as shown in Fig.~\ref{sr_fig3}(a). In this work, we set the growth rate to 16, the number of dense blocks to 3, and the number of dense layers in the $1^{st}$, $2^{nd}$, and $3^{rd}$ dense blocks are 3, 6, and 3, respectively. 

\subsection{Dataset creation} \label{s24}
To show the trained ML model's performance, careful selection of the training dataset is essential. In this paper, two different datasets are used to demonstrate our approach. First, the W2S dataset (Widefield2SIM), which includes experimentally captured diffraction-limited images (using widefield microscopy) and target images (using SIM microscopy) \cite{zhou2020w2s}. Second, the BPAE dataset, which includes experimentally captured diffraction-limited images (using custom-built multi-photon fluorescence microscopy \cite{instantFLIM_new}) and computationally generated target images (using SRRF technique \cite{SRRF_Nature}).

The W2S dataset includes 120 FOV widefield diffraction-limited fluorescence microscopy images (low-resolution: LR) and corresponding 120 FOV SIM images (high-resolution: HR). These experimental images are captured with two different fluorescence microscopy (widefield for LR images and SIM for HR images) and cells are real biological samples, namely, human cells \cite{zhou2020w2s}. In each FOV, three different channels (488nm, 561nm, and 640nm) are recorded, and we consider them as individual gray-scale images during the training and inference stages. 400 images of the same FOV are captured and averaged to generate a noise-free diffraction-limited image. Each image has a size of 512$\times$512 pixels divided into four chunks of 256$\times$256 pixels. Each FOV corresponds to \SI{51.2}{\micro\metre}$\times$\SI{51.2}{\micro\metre} (where each pixel size is 100 nm). Before the training process, all the images in the training dataset are normalized, and normalization is explained in Section.~\ref{s27}. In the case of the W2S dataset, with noise-free (average of 400 images in the same FOV) diffraction-limited images and noisy (no average of images in the same FOV) diffraction-limited images as input to the training dataset. In each case, the target image is the experimentally captured super-resolution image (SIM setup) \cite{zhou2020w2s}.

In the BPAE dataset, the BPAE sample (Invitrogen FluoCells slide \#1, F36924 contains Nuclei, F-actin, and Mitochondria) was imaged with our custom-built two-photon fluorescence microscopy system \cite{instantFLIM_new} that provides diffraction-limited images as input of the training dataset. The custom setup consists of an objective lens with 40x magnification (0.8 numerical aperture and 3.5 mm working distance). The two-photon excitation wavelength is 800 nm (for the one-photon system, the excitation wavelength is 400 nm), sample power is six mW, pixel width is 200 nm, pixel dwell-time, 12$\mathrm{\mu}$s, and the emission wavelength filter is from 300-700 nm. We used a photomultiplier tube (PMT) to convert the emission photons to current, followed by the transconductance amplifier (TA) to convert them to voltage. A total of 16 FOVs of the BPAE sample were captured, where each FOV consists of 50 diffraction-limited images, and each image has a size of 256$\times$256 pixels. The images in the 8$^{\mathrm{th}}$ FOV are used as the test dataset, and the remaining 1-7 FOVs and 9-16 FOVs data are used as the training dataset. Hence the training dataset size is 15 FOVs. We used the SRRF technique \cite{SRRF_Nature} to generate super-resolution target images from the diffraction-limited images. Fifty images of the same FOV are captured and averaged to create a noise-free diffraction-limited image. Each image has a size of 256$\times$256 pixels divided into four chunks of 128$\times$128 pixels. Before the training, all the images in the training dataset are normalized, and normalization is explained in Section.~\ref{s27}. More details of the SRRF are provided in the results section (please see Section.~\ref{sec32}). \cc{In addition, this BPAE dataset is provided as open source to validate the performance of the estimated super-resolution images when trained with small datasets. More details about the dataset are provided in the Code and Data section \ref{coderef}}. 

In this study, we show the effect of the SRDenseED method in FCNs using both W2S and the BPAE datasets.  

\subsection{Hyper-parameters} \label{s25}
Hyper-parameter search is a critical step in deep learning for quick and accurate results, primarily problem-specific and empirical. Typical hyper-parameters in FCN architecture are batch size, optimizer, and learning rate, and are carefully tuned for achieving the best fluorescence microscopy image super-resolution performance. The batch size used in the training stage is set to 3. The ``Adam" gradient descent algorithm \cc{\cite{kingma2014adam}} is used to optimize the loss function between the estimated and target super-resolution images during training. The initial learning rate is set to 3E-3, and weight decay is used to reduce the over-fitting problem to 3E-4. In addition, these parameters are fixed for all ML models: the number of feature maps in the first convolution layer is set to 48, the number of output feature maps is set to 16 ($k$-value) in every dense block, and the number of epochs is set to 400 such that the loss function reaches a stable point, the number of dense blocks to 3 and the number of dense layers in $1^{st}$, $2^{nd}$, and $3^{rd}$ dense block are 3, 6, and 3, respectively. The training time varies with the training dataset size, and for the small dataset (for 90 FOVs), the training time is less than 4 hrs on a single Nvidia 1080-ti GPU. The number of parameters (kernel weights) for simple FCN (U-Net with three encoders and three decoders) architecture and FCN with three DenseED blocks is 286,704 and 237,204, respectively. More details about the ML model architectures can be found in the Code section.

\subsection{Data processing} \label{s27}
\cc{Typically, biomedical images are too large to fit on a single GPU. Hence images are divided (input and target) into smaller patches when training the ML models. Normalization is applied as part of the pre-processing step to each image before passing it to the ML models (both simple FCNs and FCNs with the SRDenseED ML model). The input to the ML model is an image ($I$) which is linearly normalized by dividing with the maximum intensity value (here, the maximum value is 255 since images are 8-bit) and subtracting 0.5. Hence, all the pixel values passed through the ML model are always normalized ($I_{norm}$) and lie between -0.5 to 0.5 ($I_{norm} = I/255 - 0.5$). In addition, the target SR images are normalized the same as diffraction-limited images, and the pixel values lie between -0.5 to 0.5. As part of the post-processing, the output ($O_{norm}$) from the ML models is post-processed using the de-normalization step using this equation ($O_{denorm} = (O_{norm}+0.5)*255$). Finally, the estimated super-resolution images are converted to 8-bit images to match the input (DL) and target (SR) image format.} 

\subsection{Forward modeling in super-resolution imaging} \label{s28}
\cc{In the literature on computer vision or machine learning, high-resolution (HR) images are taken from a high-quality instrument which is typically expensive. Since the high-quality instrument provides minimal artifacts such as better resolution (better point spread function (PSF)) and low noise in the high-resolution images, in this case, low-resolution (LR) images are generated using forward modeling and given in $I_LR = (I_{HR} * PSF + n)$, where $I_{LR}$ is the LR image derived from HR image, $I_{HR}$ is the HR image captured using an expensive instrument, $PSF$ is the point spread function to generate LR image from HR image, $*$ is the convolution operation, $n$ additive white gaussian noise with zero mean and $\sigma$ standard deviation ~$N(0, \sigma)$. Hence, this generation method of LR images provides a blur due to the convolution of PSF, which is a 2D-Gaussian function. In this case, the ML model works as an inverse problem to detect the HR image from the LR images (which is an alternative to a conventional iterative deconvolution method \cite{fish1995blind, mannam2022deconvolution, yeh2019computational, luo2020super}). Other research areas use super-resolution in the context to upscale the low-resolution image from N$\times$N image to MN$\times$MN, where $M$ is the scaling factor, typically, $M$ is either 2, 3, or 4. Hence, the forward modeling is given by $I_{LR} = ({I_{HR}}*PSF)$, where $I_{LR}$ is the LR (down-sampled) image of size N$\times$N, $I_{HR}$ is the HR (upsampled) image of size MN$\times$MN, $PSF$ is the Gaussian function to downsample the image, and $*$ is the convolution operation. In this case, the ML model works as an inverse problem to detect the upsampled/up-scaled (HR) image from the down-sampled/down-scaled (LR) images. In contrast, in the case of optical microscopy, the low-resolution images are captured using an instrument that cannot separate close-by cells/samples \cite{prakash2022super}. Typically, this instrument is low in cost with limited resolution. Hence, the low-resolution images in this field are called ``diffraction-limited (DL) images". Also, the high-resolution images are captured using an expensive instrument/technique which provides high-resolution (which can separate the cells), and high-resolution images are called ``super-resolution (SR) images". Since both the DL and SR are captured using two different instruments, adequate data processing is required to show that both images indicate the same FOV. Hence, in our paper, the diffraction-limited and super-resolution images are from two instruments with different PSF values. Forward modeling is given as $I_{DL} = {I_{original}*PSF_{DL}}$, $I_{SR} = {I_{original}*PSF_{SR}}$ where $I_{original}$ is the true object need to image (cells or structure under a microscope), $I_{DL}$ and $I_{SR}$ are DL and SR images, respectively when the $I_{original}$ is captured with two different systems with PSF values as $PSF_{DL}$ and $PSF_{SR}$, respectively, $*$ indicates convolution operation. In this case, the ML model works as an inverse problem to detect the super-resolution (SR) images from the diffraction-limited (DL) images. For example, in the W2S dataset, the DL and SR images are captured using wide-field and SIM microscopy systems, and each instrument has a different PSF function. More details about the DL and SR images in the W2S dataset, including image acquisition systems, are provided in the original W2S paper\cite{zhou2020w2s}. Finally, in the BPAE dataset, only diffraction-limited images are captured using our custom-built FLIM system \cite{instantFLIM_new}, and corresponding super-resolution images are generated using a computation method called ``SRRF" \cite{SRRF_Nature}. More details about the BPAE datasets are provided in Section.~\ref{sec32}.}

\subsection{Evaluation metrics} \label{s26}
Several metrics are used to evaluate the estimated super-resolution images compared with the target super-resolution images. These metrics include structural similarity index measurement (SSIM)\cite{wang2004image}, peak signal-to-noise ratio (PSNR)\cite{ayas2020microscopic_fc2}, mean square error (MSE/L2 norm), mean absolute error (MAE/L1 norm), resolution scaled Pearson’s correlation coefficient (RSP)\cite{laine2019nanoj}, resolution scaled error (RSE)\cite{laine2019nanoj}, \cc{Fourier ring coefficient (FRC), which measures the close matching (in structures, brightness) of the estimated super-resolution images compared to target super-resolution images \cite{laine2019nanoj}. The smaller value of FRC indicates a better super-resolution image matching the target SR image \cite{laine2019nanoj}, with the value of 1 perfectly matching the target SR image.} The SSIM and PSNR are the most common metrics to quantify the estimation of super-resolution images \cite{ayas2020microscopic_fc2}. To quantitatively evaluate the estimated super-resolution images containing similar image features as the target super-resolution image, we calculate the structural similarity index measure (SSIM) between the two. SSIM compares luminance, brightness, and contrast values as a function of position \cite{wang2004image} and measure the similarity between two images on a scale of 0 to 1, with 1 being perfect fidelity. In addition, we evaluate the PSNR of the estimated image relative to a target super-resolution image. PSNR is the measure of mean square error (MSE) between two images normalized to the peak value in an image so that MSE between images with different bit depths or signal levels can be compared. PSNR of a given ($X$) with reference to ground truth image ($Y$) in the same FOV is defined as $\text{PSNR} (X, Y) = 10\log\left(\frac{\text{max}(Y)^2}{\text{MSE} (X, Y)}\right)$, where $\text{MSE}(X, Y) = \frac{1}{N} \sum_{n=1}^{N} (X_n - Y_n)^2 $ is the average mean-square error of $X$ and $Y$ with $N$ pixels. The highest SSIM and PSNR represent the most accurate estimation of the super-resolution image, similar to the target super-resolution image. Hence, this paper evaluates the estimated super-resolution images using SSIM and PSNR metrics. 

%%%%%%%%%%%%%%%%%%%%%%%%%%%%%%%%%%%%%%%%%%%%%%%%%%%%%%%%%%%%%%%%%%%%%%%%%%%%%
%%%%%%%%%%%%%%%%%%%%%%%%%%% Experimental Results %%%%%%%%%%%%%%%%%%%%%%%%%%%%
%%%%%%%%%%%%%%%%%%%%%%%%%%%%%%%%%%%%%%%%%%%%%%%%%%%%%%%%%%%%%%%%%%%%%%%%%%%%%
\section{Experimental results and discussion} \label{results}
%section a: write with clear images
%section b: write with noisy images
\subsection{SRDesnseED with experimental SR techniques} \label{sec31}
This section shows the training and prediction results (including 30 FOVs) with and without the SRDenseED method in FCN architecture trained using experimentally captured W2S dataset with training dataset size of 5 FOVs, 15 FOVs, 30 FOVs, 45 FOVs, 60 FOVs, 75 FOVs, and 90 FOVs. An estimated super-resolution image from the test dataset validates the trained ML model's accuracy from a diffraction-limited image during the testing phase. \cc{For the comparison purpose, we considered output from the joint denoising and super-resolution (JDSR) results from the original W2S paper \cite{zhou2020w2s} that provided the W2S dataset}. In this experiment, initially, we choose noise-free diffraction images (see Section.~\ref{s24}) that have high PSNR values as part of the training dataset since the noise in the experimental images degrades the performance of the trained ML models. Later in this section, using noisy diffraction images (see Section.~\ref{s24}) that have low PSNR values as training dataset results are illustrated. In the following experiments, a U-Net architecture \cite{UNet} with three encoder and decoder layers indicated as simple FCNs. Similarly, in the SRDesnseED method, we have selected DenseED(3,6,3) ML model as FCNs with DenseED blocks where the number of dense layers in the $1^{st}$, $2^{nd}$, and $3^{rd}$ dense blocks are 3, 6, and 3, respectively.

\subsubsection{Training performance using high PSNR W2S dataset} \label{sec311}
For the first part, the ML training dataset includes the noise-free (high PSNR) diffraction-limited images as input and SIM super-resolution images as a target, respectively. 
\begin{figure}[!ht]
\centering
\includegraphics[page=3,width=0.5\linewidth]{SRDenseED_all_figures_list.pdf}
\caption{W2S dataset average PSNR of the test dataset (includes 30 FOVs) vs. training dataset size using simple FCNs and SRDesnseED networks. Here, the ML models are trained using the high PSNR noise-free diffraction-limited images.} \label{fig_quant_noise_free_compare}
\end{figure}

First, we train a simple FCN architecture similar to the U-Net \cite{UNet} ML model, consisting of 3 encoders followed by three decoder blocks with the same small dataset. Later, we train the SRDenseED ML models with the same small dataset. The SRDenseED ML model diagram is shown in Fig.~\ref{sr_fig3}(a). Different DenseED models' performance can be checked by changing the number of dense blocks and dense layers in each dense block. We start by verifying the ML model's performance with three dense blocks but variable dense layers in each dense block. In this case, the SRDenseED method includes 3, 6, and 3 dense layers in the three dense blocks, respectively. In addition, the non-linear activation layer is set to ReLU, the loss function is the MSE loss between the estimated and target super-resolution images, the learning rate is set to 0.003, and the weight decay which is used to regularize the weights without over-fitting the model is always set to $\frac{1}{10^{\mathrm{th}}}$ of the learning rate. We perform testing on a test dataset (including 30 FOVs) of images that the model never sees during the training step. The hyperparameters are set to the same for simple FCNs and SRDenseED methods. Fig.\ref{fig_quant_noise_free_compare} shows the quantitative results of the noise-free diffraction-limited images as input and SIM images as target images in the training datasets. The SRDenseED model outperforms PSNR compared to conventional FCN networks, and this trend can be seen in the training dataset size. From Fig~\ref{fig_quant_noise_free_compare}, especially at the small training dataset size (15 FOVs), there is an average improvement of 1.35 dB in PSNR when using the SRDenseED ML model.

\begin{figure}[!ht]
\centering
\includegraphics[page=4,width=1.0\linewidth]{SRDenseED_all_figures_list.pdf}
\caption{W2S dataset PSNR (a) and SSIM (b) vs training dataset size using SRDesnseED networks trained using the high PSNR noise-free diffraction-limited images.} \label{fig_quant_noise_free_over_training}
\end{figure}

In addition, Fig.~\ref{fig_quant_noise_free_over_training} shows the quantitative results of PSNR and SSIM over the test dataset (includes 30 FOVs of 3 channels) of estimated super-resolution images from the noise-free diffraction-limited images. Based on the quantitative results of PSNR and SSIM, the SRDenseED ML models can provide better and more accurate super-resolution images than simple FCN networks when trained using a small training dataset. Even training with a small training dataset (15 FOVs) SRDenseED method can generate super-resolution images with an average PSNR improvement of 1.35 dB, and this SRDenseED method is helpful in biomedical imaging (X-ray and MRI imaging) to generate super-resolution images. In the SRDenseED method, the PSNR improvement, when trained with a 90 FOVs dataset, is only 0.71 dB more (a difference of 2.02 dB PSNR improvement from 90 FOVs and 1.31 dB from 15 FOVs training data) when compared with simple FCNs. Table~\ref{w2s_table1} shows the estimated super-resolution images' average PSNR when trained with high PSNR noise-free diffraction-limited images. Here, the SRDenseED method outperformed compared to simple FCNs when trained with a small dataset and confirmed the technique works for application-specific imaging. 

\begin{table}[!ht]
\centering
\begin{tabular}{cccc}\hline
\centering
%\minitab[c]{\shortstack{$\Delta$PSNR\\(noise-free)}}
\minitab[c]{\shortstack{Training dataset size\\(noise-free)}} & \minitab[c]{\shortstack{Simple FCN\\(PSNR (dB))}} & \minitab[c]{\shortstack{SRDenseED\\(PSNR (dB))}} & $\Delta$PSNR \\ \hline
Input & 15.78 & 15.78 & N/A \\ %\hline
5 & 19.96 & 21.27 & 1.31  \\ %\hline
15 & 20.08 & 21.43 & 1.35  \\ %\hline
30 & 19.86 & 22.37 & 2.51 \\ %\hline
45 & 20.52 & 22.47 & 1.95  \\ %\hline
60 & 20.44 & 22.51 & 2.07 \\ %\hline
75 & 20.72 & 22.99 & 2.28  \\ %\hline
90 & 20.86 & 22.89 & 2.02  \\ %\hline
\hline
\end{tabular} 
\caption{Quantitative comparison of average PSNR (dB) on test dataset (includes 30 FOVs) of simple FCNs and SRDenseED methods with different training dataset sizes. Here, the ML models are trained using the high PSNR noise-free diffraction-limited images and $\Delta$PSNR = PSNR from SRDenseED method - PSNR from simple FCNs.  }\label{w2s_table1}
\end{table}

Fig.~\ref{fig_qualitative_noise_free_T90}(a) shows one of the diffraction-limited noise-free images drawn randomly in the test dataset (10$^{th}$ FOV, channel 1) as the qualitative representation. \cc{Fig.~\ref{fig_qualitative_noise_free_T90}(b) shows the estimated super-resolution image from the pre-trained ML models given in \cite{zhou2020w2s} and is unable to show the clear structures in the estimated super-resolution image.} Fig.~\ref{fig_qualitative_noise_free_T90}(c) shows the estimated super-resolution image within the same FOV when trained with the SRDenseED ML model with a training dataset of 30 FOVs, and this image has better PSNR compared to the raw diffraction-limited image. Fig.~\ref{fig_qualitative_noise_free_T90}(d) shows the target super-resolution image captured using the SIM setup and in the same testing FOV. \cc{From Fig.~\ref{fig_qualitative_noise_free_T90}, the PSNR of the noise-free input image and estimated SR image using the JDSR method \cite{zhou2020w2s} and estimated SR image using the SRDenseED method (trained with 15 FOVs) are 19.22 dB and 17.84 dB, 22.45 dB, respectively. In this case, there is a PSNR improvement of -1.38 dB, and 3.23 dB of the randomly selected test image using the JDSR method \cite{zhou2020w2s} and our SRDenseED methods, respectively. Similarly, the SSIM values of the noise-free input image and estimated SR image using the JDSR method \cite{zhou2020w2s} and estimated SR image using the SRDenseED method (trained with 15 FOVs) are 0.64, 0.63, and 0.82, respectively. In addition, the calculated unscaled FRC value \cite{laine2019nanoj} of the noise-free input image and estimated SR image using the JDSR method \cite{zhou2020w2s} and estimated SR image using the SRDenseED method (trained with 15 FOVs) are 3.95, 4.15 and 3.77, respectively. From all quantitative metrics, our SRDenseED method provides better super-resolution images than the JDSR method.}
\begin{figure}[!ht]
\centering
\includegraphics[page=5,width=0.95\linewidth]{SRDenseED_all_figures_list.pdf}
\caption{Sample from the W2S dataset (a) noise-free diffraction-limited image, \cc{(b) estimated super-resolution image from JDSR method \cite{zhou2020w2s},} (c) estimated super-resolution image from the SRDenseED \cc{(Ours) ML model} (test image is taken from 10$^{th}$FOV, channel 1) and (d) is the experimentally captured target super-resolution using SIM microscopy. Here the input sample is a diffraction-limited noise-free image. The top row indicates the full frame (of size 512 $\times$ 512), and the bottom row indicates the region of interest (ROI: marked in the yellow square of size 100 $\times$ 100) from the respective top row images. Scale bar: 10 $\mu$m.} \label{fig_qualitative_noise_free_T90}
\end{figure}
%noisy section
\subsubsection{Training performance using low PSNR W2S dataset} \label{sec312}
However, obtaining noise-free images in real-time measurements is difficult (when dynamic processes are included) and time-consuming (needing multiple averages with the same FOV). Hence, the following results show the performance of our demonstrated SRDenseED ML model when trained on diffraction-limited noisy images.
\begin{figure}[!ht]
\centering
\includegraphics[page=6,width=0.5\linewidth]{SRDenseED_all_figures_list.pdf}
\caption{W2S dataset average PSNR of the test dataset (includes 30 FOVs) vs. training dataset size using simple FCNs and SRDesnseED networks. Here, the ML models are trained using the low PSNR noisy diffraction-limited images.} \label{fig_quant_noisy_compare}
\end{figure}
The response of the trained ML models using a small dataset with simple FCNs and SRDenseED ML models are analyzed with noisy diffraction-limited images as input. Fig.\ref{fig_quant_noisy_compare} shows the quantitative results of the noisy diffraction-limited images as input and SIM images as target images in the training datasets. The SRDenseED model outperforms PSNR compared to simple FCNs, and this trend can be seen over the training dataset size (even though the images are noisy and diffraction-limited). From Fig.~\ref{fig_quant_noisy_compare}, especially at the small training dataset size (15 FOVs), there is an average improvement of 0.92 dB in PSNR when using the SRDenseED ML model. In addition, Fig.~\ref{fig_quant_noisy_over_training} shows the quantitative results of PSNR and SSIM over the test dataset (includes 30 FOVs of 3 channels). Based on the quantitative results of PSNR and SSIM, the SRDenseED ML models can provide better and more accurate super-resolution images when trained with a small training dataset. Table~\ref{w2s_table2} shows the estimated super-resolution image quantitative metrics when trained with low PSNR noisy diffraction-limited images. Again, the SRDenseED method outperformed compared to simple FCNs when trained with a small dataset and confirmed the technique works for application-specific imaging. Here the results are not meant to be used as any generalized super-resolution images instead the results are meant for the application-specific imaging modalities/configurations. 
\begin{figure}[!ht]
\centering
\includegraphics[page=7,width=1.0\linewidth]{SRDenseED_all_figures_list.pdf}
\caption{W2S dataset PSNR (a) and SSIM (b) vs. training dataset size using SRDesnseED networks trained using the low PSNR noisy diffraction-limited images.} \label{fig_quant_noisy_over_training}
\end{figure}

\begin{figure}[!ht]
\centering
\includegraphics[page=8,width=0.95\linewidth]{SRDenseED_all_figures_list.pdf}
\caption{Sample from the W2S dataset (a) noisy diffraction-limited image, \cc{(b) estimated super-resolution image from JDSR method \cite{zhou2020w2s},} (c) estimated super-resolution image from the SRDenseED (Ours) ML model (test image is taken from 10$^{th}$FOV, channel 1), and (d) is the experimentally captured target super-resolution using SIM microscopy. Here the input sample is a diffraction-limited noisy image. The top row indicates the full frame (of size 512 $\times$ 512), and the bottom row indicates the region of interest (ROI: marked in the yellow square of size 100 $\times$ 100) from the respective top row images. Scale bar: 10 $\mu$m.} \label{fig_qualitative_noisy_T90}
\end{figure}

\begin{table}[!ht]
\centering
\begin{tabular}{cccc}\hline
\centering
%\minitab[c]{\shortstack{$\Delta$PSNR\\(noise-free)}}
\minitab[c]{\shortstack{Training dataset size\\(noisy)}} & \minitab[c]{\shortstack{Simple FCN\\(PSNR (dB))}} & \minitab[c]{\shortstack{SRDenseED\\(PSNR (dB))}} & $\Delta$PSNR \\ \hline
Input & 15.67 & 15.67 & N/A \\ %\hline
5 & 19.54 & 19.86 & 0.31  \\ %\hline
15 & 19.45 & 20.38 & 0.92  \\ %\hline
30 & 19.86 & 20.78 & 0.91 \\ %\hline
45 & 20.09 & 20.76 & 0.68  \\ %\hline
60 & 20.04 & 20.72 & 0.68 \\ %\hline
75 & 20.52 & 20.84 & 0.32  \\ %\hline
90 & 20.19 & 21.20 & 1.01  \\ %\hline
\hline
\end{tabular} 
\caption{Quantitative comparison of average PSNR (dB) on test dataset (includes 30 FOVs)  of simple FCNs and SRDenseED methods trained with different noisy dataset sizes. Here, the ML models are trained using the low PSNR noisy diffraction-limited images and $\Delta$PSNR = PSNR from SRDenseED method - PSNR from simple FCNs.}\label{w2s_table2}
\end{table}
Fig.~\ref{fig_qualitative_noisy_T90}(a) shows one of the diffraction-limited noisy images in a test dataset (10$^{th}$ FOV, channel 1). \cc{Fig.~\ref{fig_qualitative_noisy_T90}(b) shows the estimated super-resolution image from the pre-trained ML models given in \cite{zhou2020w2s} and is unable to show the clear structures in the estimated super-resolution image.} Fig.~\ref{fig_qualitative_noisy_T90}(c) shows the estimated super-resolution image within the same FOV when trained with the SRDenseED ML model, and this image has better PSNR compared to the raw diffraction-limited image. Fig.~\ref{fig_qualitative_noisy_T90}(d) shows the target super-resolution image captured by SIM setup within the same testing FOV. \cc{From Fig.~\ref{fig_qualitative_noisy_T90}, the PSNR of the noisy input image and estimated SR image using the JDSR method in W2S paper, and the estimated SR image using the SRDenseED method (trained with 15 FOVs) are 16.72 dB and 17.41 dB, 20.11 dB, respectively. Hence, in this case, a PSNR improvement of 0.69 dB and 3.39 dB of the randomly selected test image using the JDSR method and our SRDenseED methods, respectively. Similarly, the SSIM values of the noisy input image and estimated SR image using the JDSR method and estimated SR image using the SRDenseED method (trained with 15 FOVs) are 0.19, 0.59, and 0.69, respectively. As expected, the JDSR method improved PSNR when the input image is noisy compared to noise-free, where the significant contribution is from the image denoising step. In addition, the calculated unscaled FRC value \cite{laine2019nanoj} of the noise-free input image and estimated SR image using the JDSR method \cite{zhou2020w2s} and estimated SR image using the SRDenseED method (trained with 15 FOVs) are 5.80, 5.59 and 5.43, respectively. From all quantitative metrics, our SRDenseED method provides better super-resolution images than the JDSR method.} We observe that our SRDenseED method (trained with 15 FOVs) provides accurate super-resolution images by providing an average PSNR improvement of 5.65 (21.43-15.78, see Table.~\ref{w2s_table1}) dB and 4.71 (20.38-15.67, see Table.~\ref{w2s_table2}) dB in noise-free diffraction-limited images as input and noisy diffraction-limited images as input, respectively. In addition, compared to simple FCN architecture, our SRDenseED method (trained with 15 FOVs) provided an average PSNR improvement of 1.35 dB and 0.92 dB, in the case of noise-free and noisy diffraction-limited input images, respectively. 

\subsection{SRDesnseED with computational SR techniques} \label{sec32}
Generating super-resolution images requires an additional experimental setup, which is expensive, and the research labs may not have this setup. However, experimental diffraction-limited image generation is a typical setup, and super-resolution images can be generated using computational methods. For example, super-resolution radial fluctuation (SRRF) \cite{SRRF_Nature} is a computational method to generate super-resolution images within the same FOV from multiple diffraction-limited images (captured with different time instances). In this section, we captured experimental diffraction-limited images of BPAE samples (Invitrogen FluoCells slide\#1 F36924, mitochondria labeled with MitoTracker Red CMXRos, F-actin labeled with Alexa Fluor 488 phalloidin, nuclei labeled with DAPI) using our custom-built two-photon fluorescence microscopy system \cite{instantFLIM_new}. In this step, the captured images include noise. The custom setup consists of an objective lens with 40x magnification (0.8 numerical aperture and 3.5 mm working distance). The two-photon excitation wavelength is 800 nm (for the one-photon system, the excitation wavelength is 400 nm), sample power is six mW, pixel width is 200 nm, pixel dwell-time, 12$\mathrm{\mu}$s, and the emission wavelength filter is from 300-700 nm. In our imaging system, all the fluorophores-labeled organelles are excited together using a single excitation wavelength (in this case, 800 nm) and get the collective emission together using a bandpass filter (300-700 nm) that shows all the fluorophores together in the fluorescence intensity image. We used a photomultiplier tube (PMT) to convert the emission photons to current, followed by the transconductance amplifier (TA) to convert them to voltage. More details about the setup can be found in \cite{instantFLIM_new}. A total of 16 different FOVs (small training dataset) of the BPAE sample are captured using our system, where each FOV consists of 50 raw images, and each image has a size of 256$\times$256. The target super-resolution images are generated using the SRRF technique. SRRF method performs two steps \cite{SRRF_Nature}, i.e., spatial and temporal steps, to generate super-resolution images. Spatial SRRF estimates and maps the most likely positions of the molecules, followed by temporal SRRF to improve the resolution of the final super-resolution SRRF image using spatial resolution step statistics. The center of the fluorophores is estimated and mapped to a ``radiality" map in simple terms. SRRF method provides the super-resolution image in the sub-pixel range (with a magnification of 5 times by default) and reshapes it (using bilinear interpolation) to the raw image dimension 256$\times$256. Note: SRRF can provide inaccurate target results if the parameters are not set correctly during this target generation stage, and more details can be found in \cite{SRRF_Nature}.

The experimentally captured diffraction-limited images (also noisy) and SRRF-generated images are used as the input and target of the small training dataset, respectively. Normalization is applied to each image before passing it to the FCNs with the SRDenseED ML model. The image normalization is conducted by dividing the maximum value in the data type (here, the maximum value is 255) and subtracting 0.5. Hence, all the pixel values passed through the ML model are always normalized and lie between -0.5 to 0.5. The images generated in the 8$^{\mathrm{th}}$ FOV are used as the test dataset, and the remaining 1-7 FOVs and 9-16 FOVs data are used as the training dataset. The training dataset consists of 15 FOVs, called a ``small training dataset". Here the input is a 16-bit grayscale channel. 

\begin{figure}[!ht]
\centering
\includegraphics[page=9,width=1.0\linewidth]{SRDenseED_all_figures_list.pdf}
\caption{BPAE sample diffraction-limited image (a) acquired with our custom-built two-photon microscope \cite{instantFLIM_new}, noise-free image (averaged within the same FOV) (b) and the target super-resolution image generated by SRRF method (c). Estimated super-resolution image using the trained ML model with dense blocks in FCN (d). The resolution improvement in (e) includes the line plots of images shown in (a, b, c, and d in blue color), respectively, with markers in wine and blue colors indicating the full width half maximum (FWHM) of diffraction-limited and estimated super-resolution images. The top row (a, b, c, and d) indicates the full frame (of size 256 $\times$ 256), and the bottom row (f, g, h, and i) indicates the region of interest (ROI: marked in the green square of size 75 $\times$ 75) from the respective top row images. Pixel width, 200nm; pixel dwell-time, 12$\mathrm{\mu}$s; excitation power, 6 mW.} \label{fig_main}
\end{figure}

The quantitative and qualitative results from the test dataset are shown in Fig.~\ref{fig_main} after training the ML model using the SRDesnseED method. Fig.~\ref{fig_main}(a) shows the experimentally captured (using a custom two-photon FLIM system) nosy diffraction-limited image of the BPAE sample cell, and Fig.~\ref{fig_main}(b) indicates a noise-free diffraction-limited image within the same FOV. Similarly, Fig.~\ref{fig_main}(a) shows the target super-resolution image generated using the computation SRRF method from multiple diffraction-limited images. Fig.~\ref{fig_main}(d) shows the estimated super-resolution images from the DenseED (3,6,3) configuration ML model. The estimated super-resolution image accurately estimates sub-micron features (mitochondria) and is comparable with the target image. Averaging more images within the same FOV improves the PSNR (from 21.24 dB to 21.89 dB), but unable to find the sub-micron super-resolution structures (see Fig.~\ref{fig_main}(b)). The PSNR values of the noisy diffraction-limited image, noise-free diffraction-limited image, and the estimated SRDenseED image are 21.24 dB, 21.89 dB, and 24.73 dB, as shown in Fig.~\ref{fig_main}(a), (b), (d) respectively (with respect to target image as shown in Fig.~\ref{fig_main}(c)). Hence, there is a 3.49 dB improvement in PSNR from the trained SRDenseED method compared to the diffraction-limited noisy test image. The improvement in the PSNR is due to the identification of small features, and the estimated image closely matches the target image. Hence, the trained ML model with the SRDenseED method can achieve super-resolution from the diffraction-limited images even though the training dataset size is limited. In addition, Fig.~\ref{fig_main}(e) provides the qualitative and quantitative metrics on the estimated super-resolution image with a marked region and corresponding line plots of the trained ML model using the DenseED model with three dense blocks and 3,6,3 are the dense layers in each dense block respectively. The FWHM for the diffraction-limited and estimated super-resolution images are $\approx$ \SI{1.2}{\micro\metre} and $\approx$ \SI{0.6}{\micro\metre}, respectively, which shows at least 2$\times$ resolution improvement. The top row in Fig.~\ref{fig_main}(a, b, c, and d) indicates the full frame (of size 256 $\times$ 256), and the bottom row in Fig.~\ref{fig_main}(f, g, h, and i) indicates the region of interest (ROI: marked in the green square of size 75 $\times$ 75) from the respective full-FOV images. 

Additional qualitative and quantitative results with different DenseED configurations are provided in the GitHub repository (\url{https://github.com/ND-HowardGroup/Application-Specific-Super-resolution.git}) on the estimated super-resolution images of the trained ML models. Variations of the estimated SR images PSNR and SSIM are shown, including variation in the learning rate, non-linear activation function, sample dataset size, and the loss function as the mixed loss of MSE loss and SSIM loss to optimize the MSE loss and SSIM loss simultaneously in FCN architecture. Also, these demonstrated DenseED blocks could be applied to estimate super-resolution images from resolution-limited images with GAN architecture with retraining (more results are shown in the GitHub repository for the W2S dataset and BPAE dataset). 

\cc{If the test dataset is entirely different from the training dataset, generated super-resolution images might have some artifacts in the output \cite{wang2018esrgan}. Also, if the target generation has some artifacts, then the estimated super-resolution using this trained dataset will also have artifacts. Consider the BPAE dataset, where the target image is generated using the SRRF computational method, which can provide super-resolution images with artifacts if the computational parameters are not appropriately set \cite{SRRF_Nature}. In this case, the inaccuracy of the ground truth image will affect the performance of the ML model. In addition, the generalization capability of the trained ML model is limited when trained using a small training dataset might also include artifacts such as hallucination effects, blur, and other cells to display where the estimated super-resolution image has more details than the ground-truth super-resolution image. Hence it is always recommended to check if the generated super-resolution images have any hallucinations or artifacts using the existing quantitative metrics such as PSNR, SSIM, and FRC, as mentioned in Section.~\ref{s26}. To reduce artifacts, additional steps are required when generating super-resolution images, such as using residual layers \cite{nasrollahi2020deep}. }   

Finally, the DenseED block in ML model architectures helps to generate super-resolution images when the ML model is trained with a small dataset. The performance improvement depends on optimizing other hyper-parameters and parameters of the network, including learning rate, non-linear activation, loss function, and weight decay, on regularizing the over-fitting. For the SRDenseED method, the number of dense blocks and dense layers are also significant in each dense block. Clearly, from the above experiments, the SRDenseED method provides accurate results compared to simple FCNs.  

%%%%%%%%%%%%%%%%%%%%%%%%%%%%%%%%%%%%%%%%%%%%%%%%%%%%%%%%%%%%%%%%%%%%%%%%%%%%%
%%%%%%%%%%%%%%%%%%% Discussion and  Conclusion %%%%%%%%%%%%%%%%%%%%%%%%%%%%%%
%%%%%%%%%%%%%%%%%%%%%%%%%%%%%%%%%%%%%%%%%%%%%%%%%%%%%%%%%%%%%%%%%%%%%%%%%%%%%
\section{Conclusions} \label{conclusions} %1page here 
Machine learning models have been previously demonstrated to generate super-resolution from diffraction-limited images. Such approaches require thousands of training images, which is prohibitively difficult in many biological samples. We showed the FCN architectures with the SRDenseED method, including Dense Encoder-Decoder blocks, to train super-resolution FCNs using a small training dataset. Our results show an accurate estimation of super-resolution images with denseED blocks in conventional ML models (see Fig.~\ref{fig_qualitative_noise_free_T90}(b), Fig.~\ref{fig_qualitative_noisy_T90}(b) and Fig.~\ref{fig_main}(d)). We showed the estimated super-resolution image PSNR results and compared them with the target super-resolution images in the case of both experimentally captured SIM setup (as shown in Section.~\ref{sec31}) and computationally generated with the SRRF method (as shown in Section.~\ref{sec32}), with PSNR improvement of 3.66 dB (in case of noise-free DL images) and 3.49 dB, respectively. \cc{Our primary focus was to demonstrate the new ML method (our SRDenseED method) capable of providing application-specific super-resolution (for example, fluorescence microscopy) images when trained using a small training dataset.} In addition, we used the SRRF method for the target generation since it is computational and easy to use. Besides, our demonstrated model can work with other super-resolution target generation methods like STED/STORM/PALM/SIM. While we evaluated the technique on super-resolution fluorescence microscopy, this approach shows promise for an extension to other deep learning based image enhancements (e.g., image denoising networks \cite{weigert2018content, mannam2022real}, image super-resolution \cite{li2018dlbi, zhao2022sparse, kaderuppan2022net, zelger2018three, mannam2021deep}, image segmentation networks \cite{UNet}, and other imaging modalities like X-ray \cite{hemdan2020covidx, jain2021deep, baltruschat2019comparison} and MRI imaging \cite{liu2018applications}).

\section*{Disclosures}
\noindent The authors declare no conflicts of interest.

\section*{Funding}
This material is based upon work supported by the National Science Foundation (NSF) under Grant No. CBET-1554516 and the National Science Foundation (NSF) funding program named Industry-University Cooperative Research Program (IUCRC) Center for Bioanalytical Metrology (CBM) under Grant No. 1916601. 
%The authors are grateful for funding this work. 

\section*{Acknowledgments} %\acknowledgments % equivalent to \section*{ACKNOWLEDGMENTS}       
The authors further acknowledge the Notre Dame Center for Research Computing (CRC) for providing the Nvidia GeForce GTX 1080-Ti GPU resources for training the Super-Resolution (SR) Fluorescence Microscopy dataset in Pytorch.

\section*{Code and Data} \label{coderef}
The results used in this manuscript are open-source and can be accessed via GitHub provided at \url{https://github.com/ND-HowardGroup/Application-Specific-Super-resolution.git}. The code and other resources are provided for public access to obtain the results reported in the manuscript. The Super-Resolution (SR) Fluorescence Microscopy dataset, including the diffraction-limited and super-resolution images used to train the DenseED ML models, are provided in open-source \url{https://curate.nd.edu/show/5h73pv66g4s} and my Ph.D. thesis \cite{mannam2022overcoming}. 

% References
\bibliography{report} % bibliography data in report.bib
\bibliographystyle{spiebib} % makes bibtex use spiebib.bst

\section*{Biography} %add biography of each author

Varun Mannam was born in Ongole, Andhra Pradesh, India, and received a Bachelor of Technology and Master of Technology degree in Electronics and Communications Engineering from the Bapatla Engineering College, Acharya Nagarjuna University, Guntur, Andhra Pradesh, India, and Indian Institute of Technology Kharagpur (IIT Kharagpur), West Bengal, India, in 2010 and 2012, respectively. From 2012-2017, he worked as Staff Software Engineer to develop wireless connectivity communication protocols (such as Bluetooth and WIFI) at National Instruments R$\&$D Bangalore, India. He came to the University of Notre Dame in 2017 to pursue a doctorate in Electrical Engineering. Currently, he is a Ph.D. student working with Dr. Scott Howard in the Biophotonics Research Group. His research is about the applications of fluorescence microscopy image enhancements that could advance the work of other researchers and medical personnel in various fields. His research interests include deep learning, machine learning, computer vision, FLIM, super-resolution, compressive sensing, and neural circuits. 

Prof. Scott Howard received his Ph.D. degree from the Department of Electrical Engineering, Princeton University, Princeton, NJ, in 2008. His research at Princeton University focused on high-performance quantum cascade laser design, fabrication, and characterization. Scott Howard was the 2007 Newport Award winner for excellence in photonics research at Princeton University. He was a Postdoctoral Research Associate with the School of Applied and Engineering Physics, Cornell University, Ithaca, NY, where his research focused on fiber-based nonlinear medical imaging. He is currently an associate professor working at the University of Notre Dame. Prof. Scott Howard’s research focuses on how the interaction of photons and tissue can be used to aid diagnosis and fundamental research in biological fields. The group’s work includes developing new techniques to image molecules in real-time, in living tissue in 3D (e.g., fast, full frame, super-resolution FLIM), contrast agent development (e.g., encapsulation of nonlinear optical dyes), and open-source software tools to enhance biomedical research’s ability to produce low-noise, super-resolution microscopy images using conventional laboratory microscopes.

\pagebreak

\section*{Figure captions} 
\begin{itemize}
\item Figure 1: Block diagram of fully convolutional networks with skip connections, including the encoder and decoder blocks. Here the network indicates the autoencoder (AE) and U-Net architectures without and with skip connections \cite{mannam2020instant}, respectively. Encoder and decoder blocks consist of batch-norm, ReLU (rectified linear unit), and convolution layers. Conv(s2) and Conv$^{T}$(s2) indicate the convolution and convolution transpose layers with a stride of 2, respectively. The symbol $\oplus$ represents the concatenation layer that combines the output from the encoder layer and decoder layer in the number of channels dimension.
\item Figure 2: Block diagram of Fully convolutional networks with dense blocks (a). dense blocks consist of multiple dense layers (b) where each dense layer's input feature maps are concatenated progressively. The dense layer (c) consists of the batch norm, ReLU, and convolution layer with stride 1 in sequence order.
\item Figure 3: W2S dataset average PSNR of the test dataset (includes 30 FOVs) vs. training dataset size using simple FCNs and SRDesnseED networks. Here, the ML models are trained using the high PSNR noise-free diffraction-limited images.
\item Figure 4: W2S dataset PSNR (a) and SSIM (b) vs training dataset size using SRDesnseED networks trained using the high PSNR noise-free diffraction-limited images.
\item Figure 5: Sample from the W2S dataset (a) noise-free diffraction-limited image, (b) estimated super-resolution image from JDSR method \cite{zhou2020w2s}, (c) estimated super-resolution image from the SRDenseED (Ours) ML model (test image is taken from 10$^{th}$FOV, channel 1) and (d) is the experimentally captured target super-resolution using SIM microscopy. Here the input sample is a diffraction-limited noise-free image. The top row indicates the full frame (of size 512 $\times$ 512), and the bottom row indicates the region of interest (ROI: marked in the yellow square of size 100 $\times$ 100) from the respective top row images. Scale bar: 10 $\mu$m.
\item Figure 6: W2S dataset average PSNR of the test dataset (includes 30 FOVs) vs. training dataset size using simple FCNs and SRDesnseED networks. Here, the ML models are trained using the low PSNR noisy diffraction-limited images.
\item Figure 7: W2S dataset PSNR (a) and SSIM (b) vs. training dataset size using SRDesnseED networks trained using the low PSNR noisy diffraction-limited images.
\item Figure 8: Sample from the W2S dataset (a) noisy diffraction-limited image, (b) estimated super-resolution image from JDSR method \cite{zhou2020w2s}, (c) estimated super-resolution image from the SRDenseED (Ours) ML model (test image is taken from 10$^{th}$FOV, channel 1), and (d) is the experimentally captured target super-resolution using SIM microscopy. Here the input sample is a diffraction-limited noisy image. The top row indicates the full frame (of size 512 $\times$ 512), and the bottom row indicates the region of interest (ROI: marked in the yellow square of size 100 $\times$ 100) from the respective top row images. Scale bar: 10 $\mu$m.
\item Figure 9: BPAE sample diffraction-limited image (a) acquired with our custom-built two-photon microscope \cite{instantFLIM_new}, noise-free image (averaged within the same FOV) (b) and the target super-resolution image generated by SRRF method (c). Estimated super-resolution image using the trained ML model with dense blocks in FCN (d). The resolution improvement in (e) includes the line plots of images shown in (a, b, c, and d in blue color), respectively, with markers in wine and blue colors indicating the full width half maximum (FWHM) of diffraction-limited and estimated super-resolution images. The top row (a, b, c, and d) indicates the full frame (of size 256 $\times$ 256), and the bottom row (f, g, h, and i) indicates the region of interest (ROI: marked in the green square of size 75 $\times$ 75) from the respective top row images. Pixel width, 200nm; pixel dwell-time, 12$\mathrm{\mu}$s; excitation power, 6 mW.
\end{itemize}
\end{document} 
