## Supplementary figures and images for "Small Training Dataset Convolutional Neural Networks for Application Specific Super-Resolution Microscopy"

### SRDenseED_all_figures_list.pdf

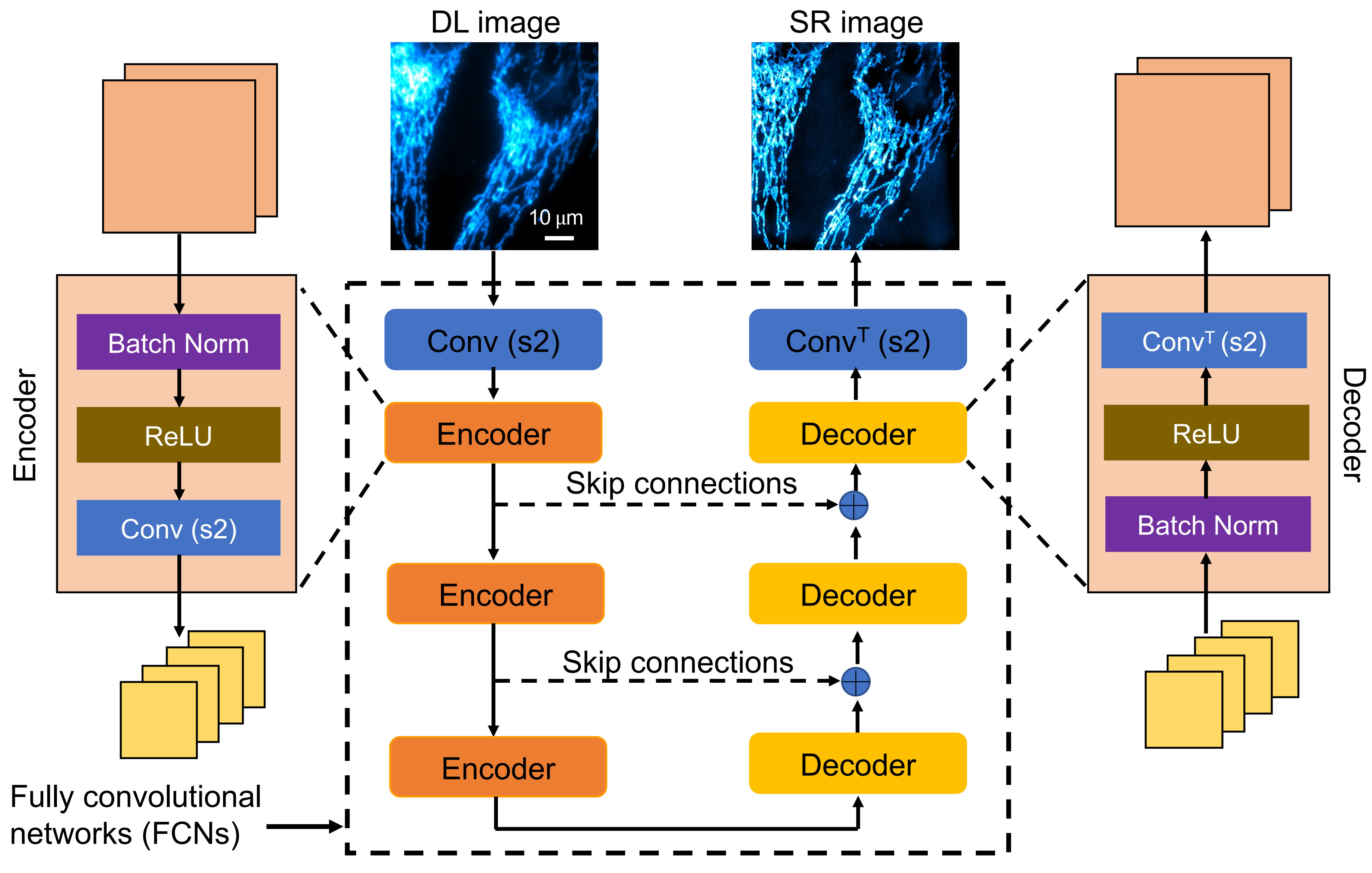

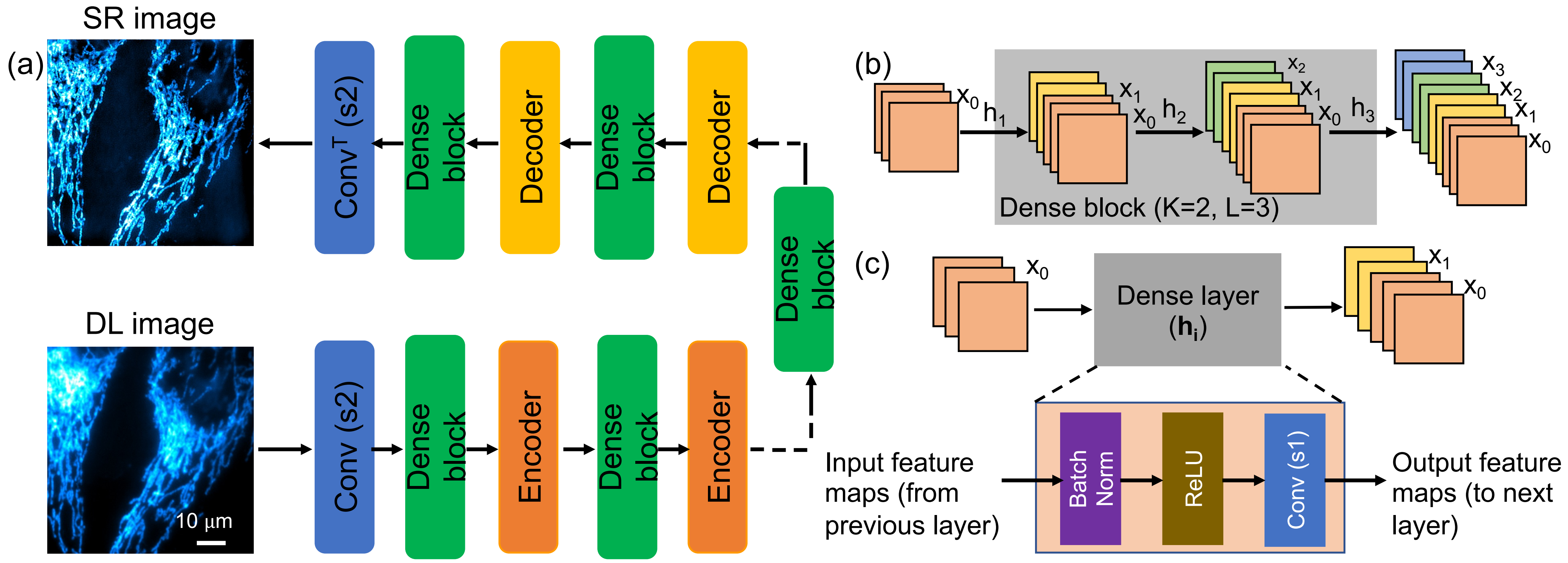

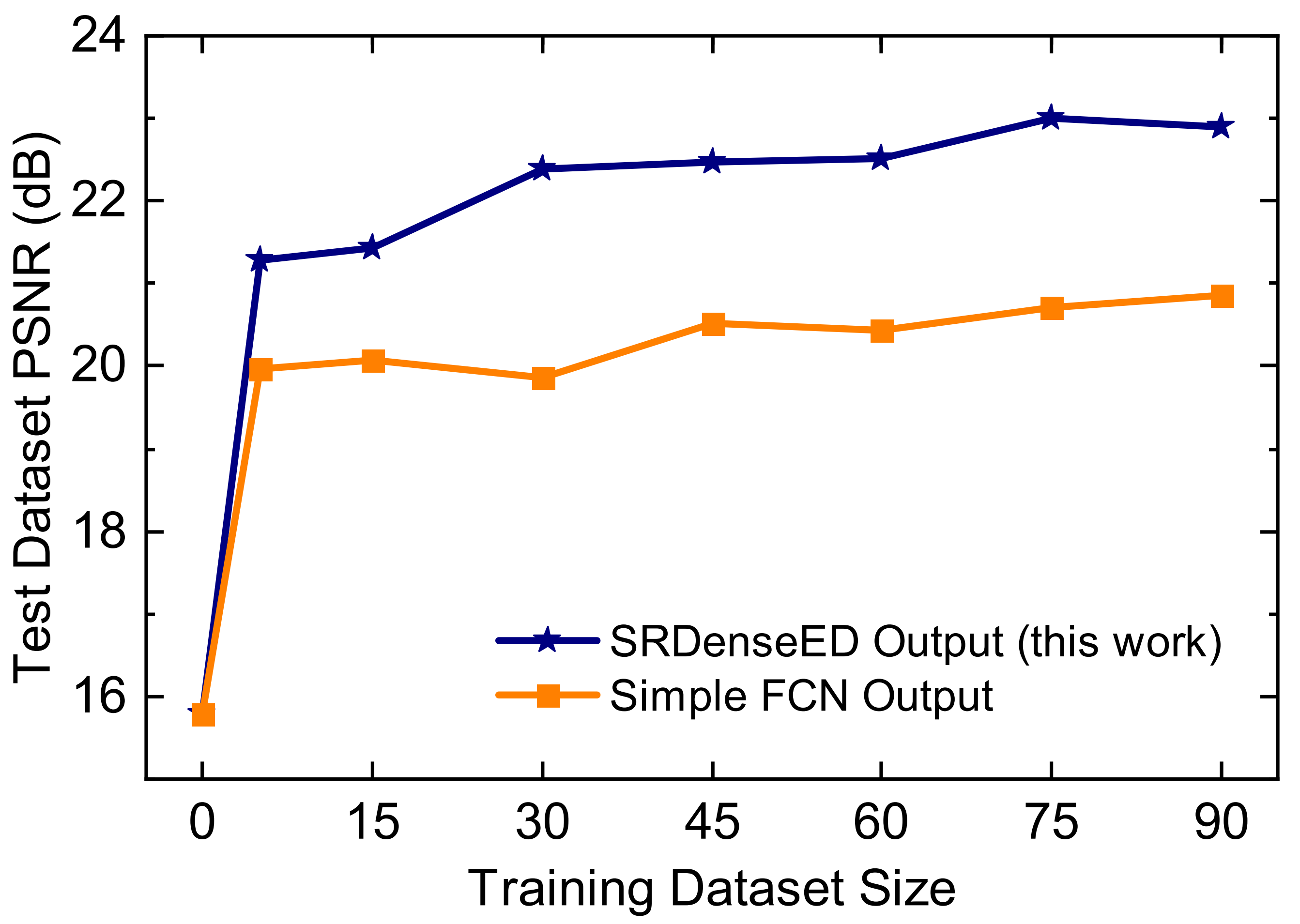

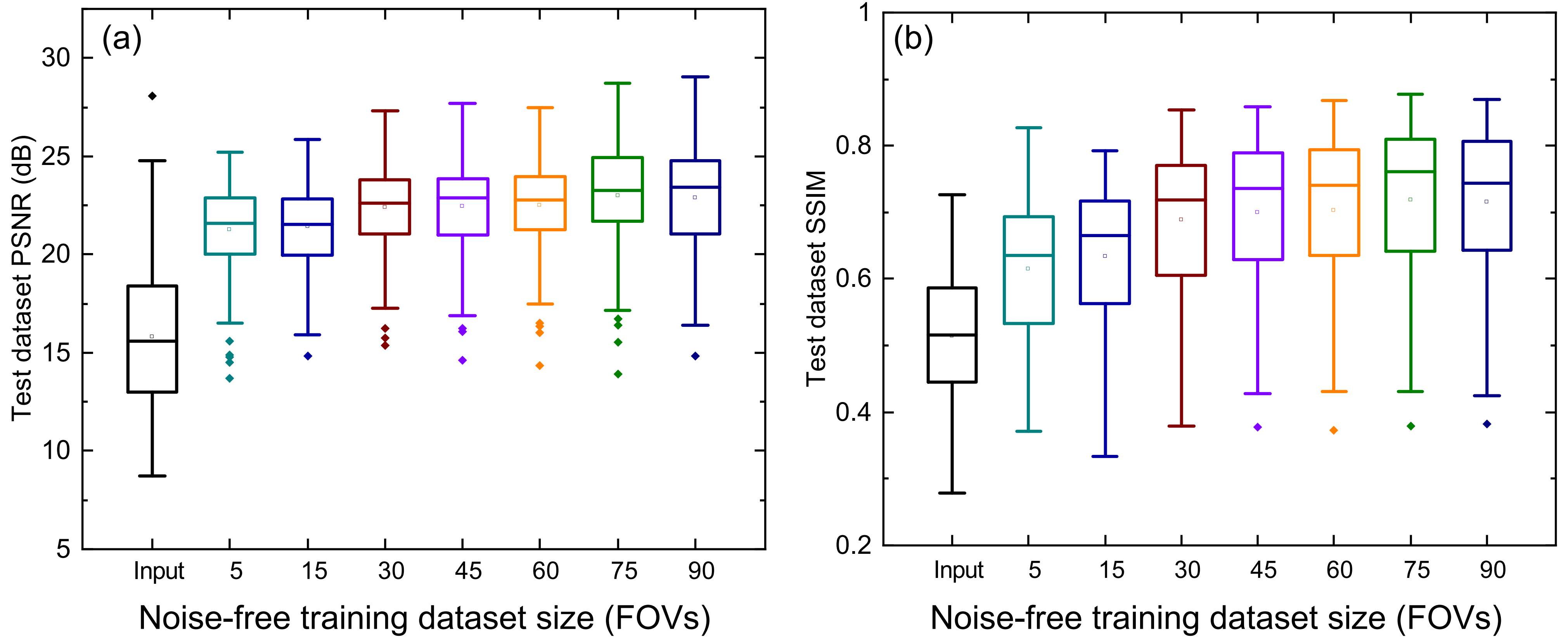

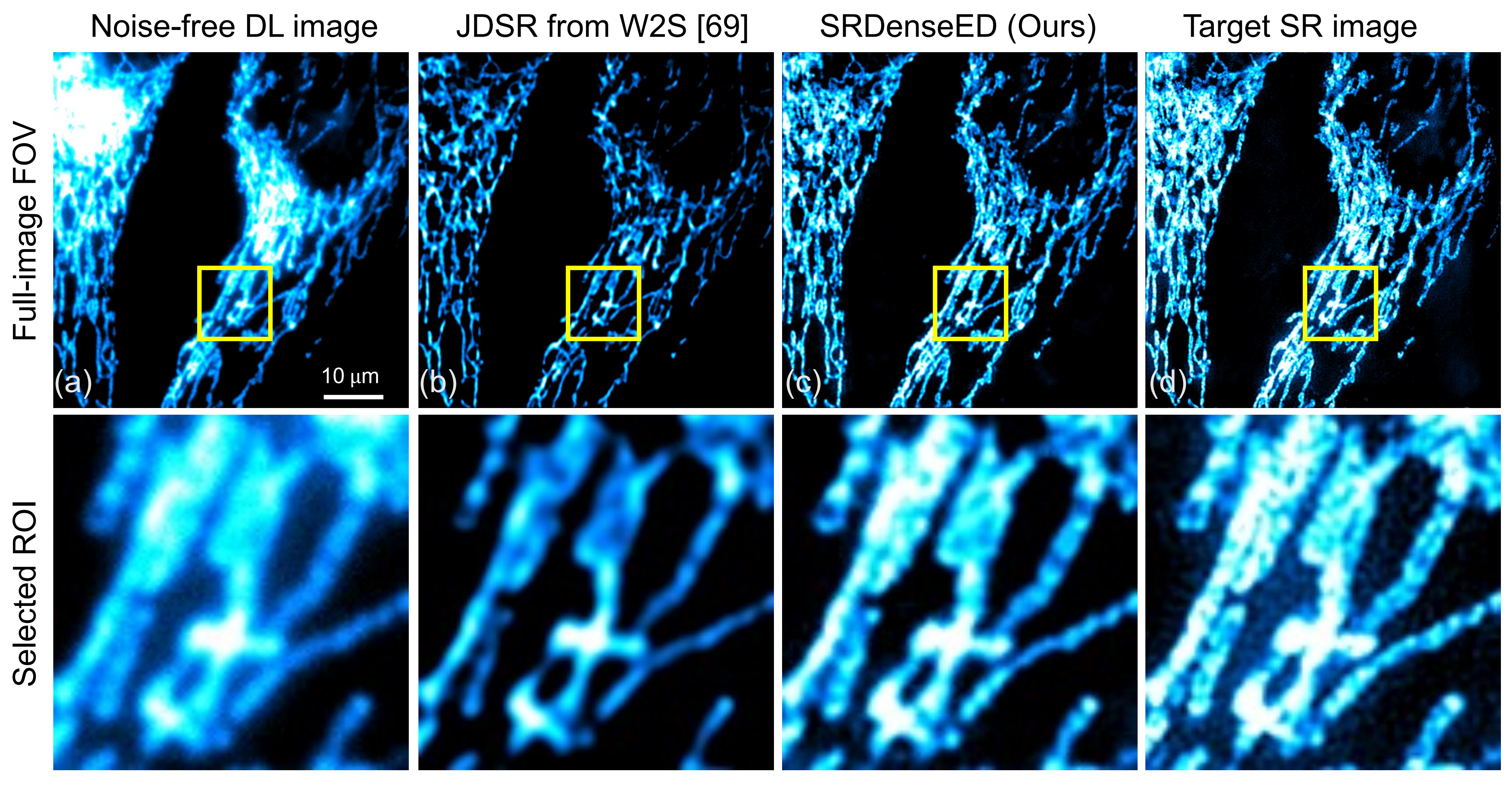

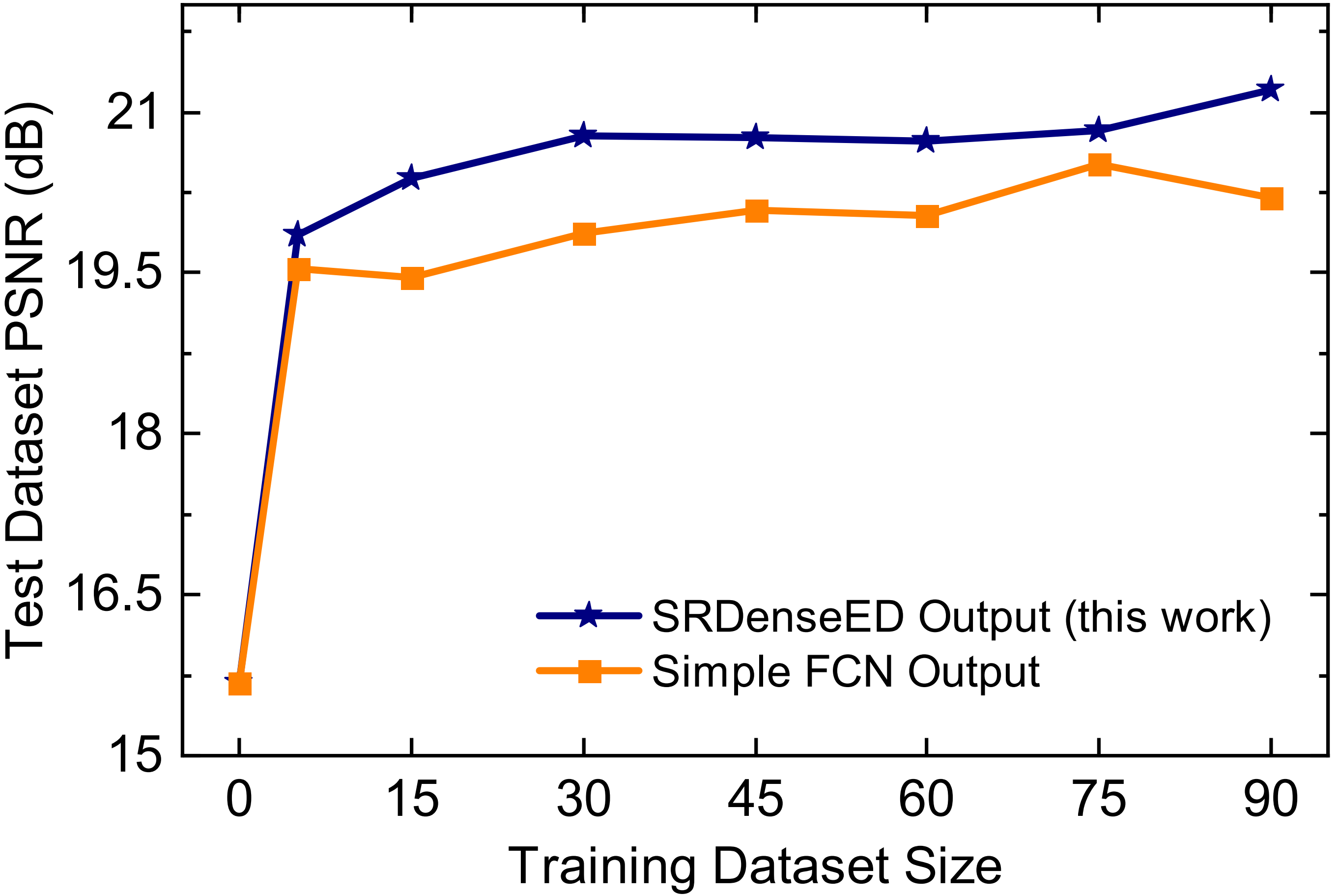

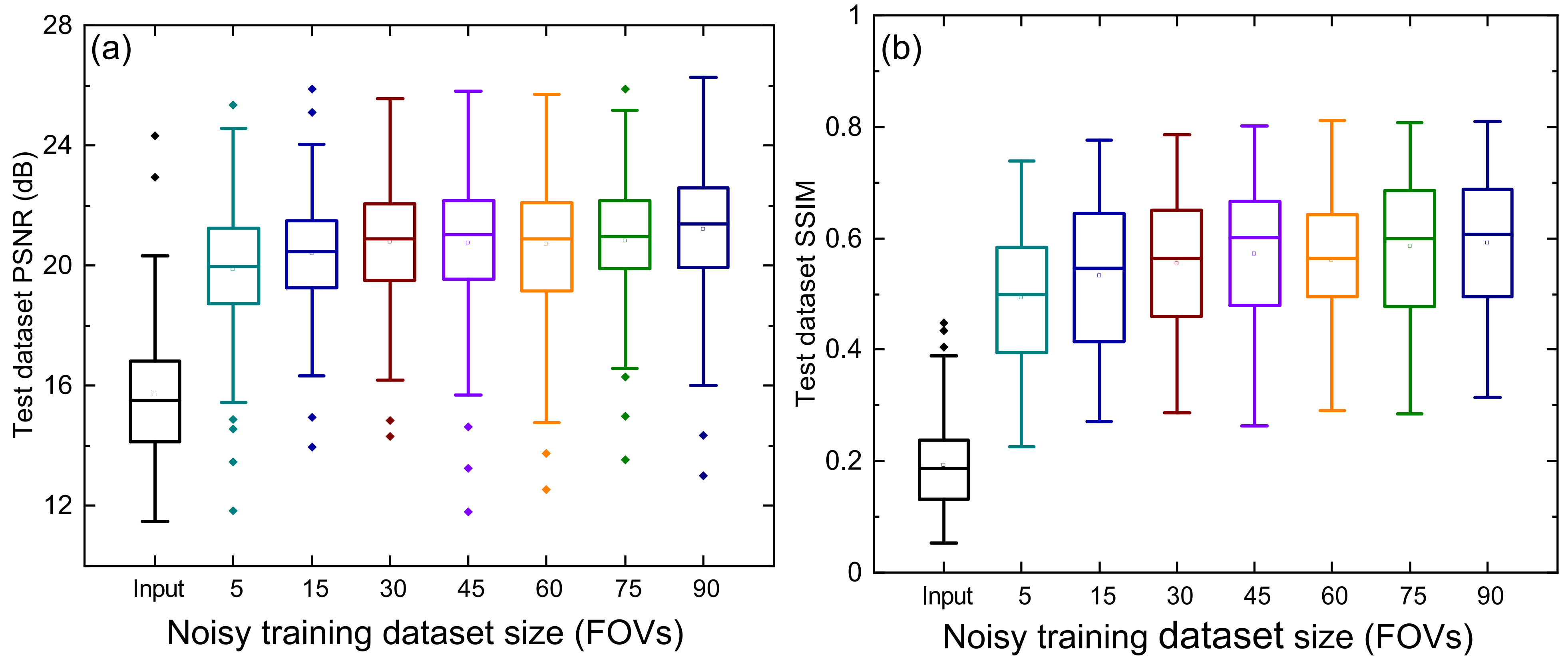

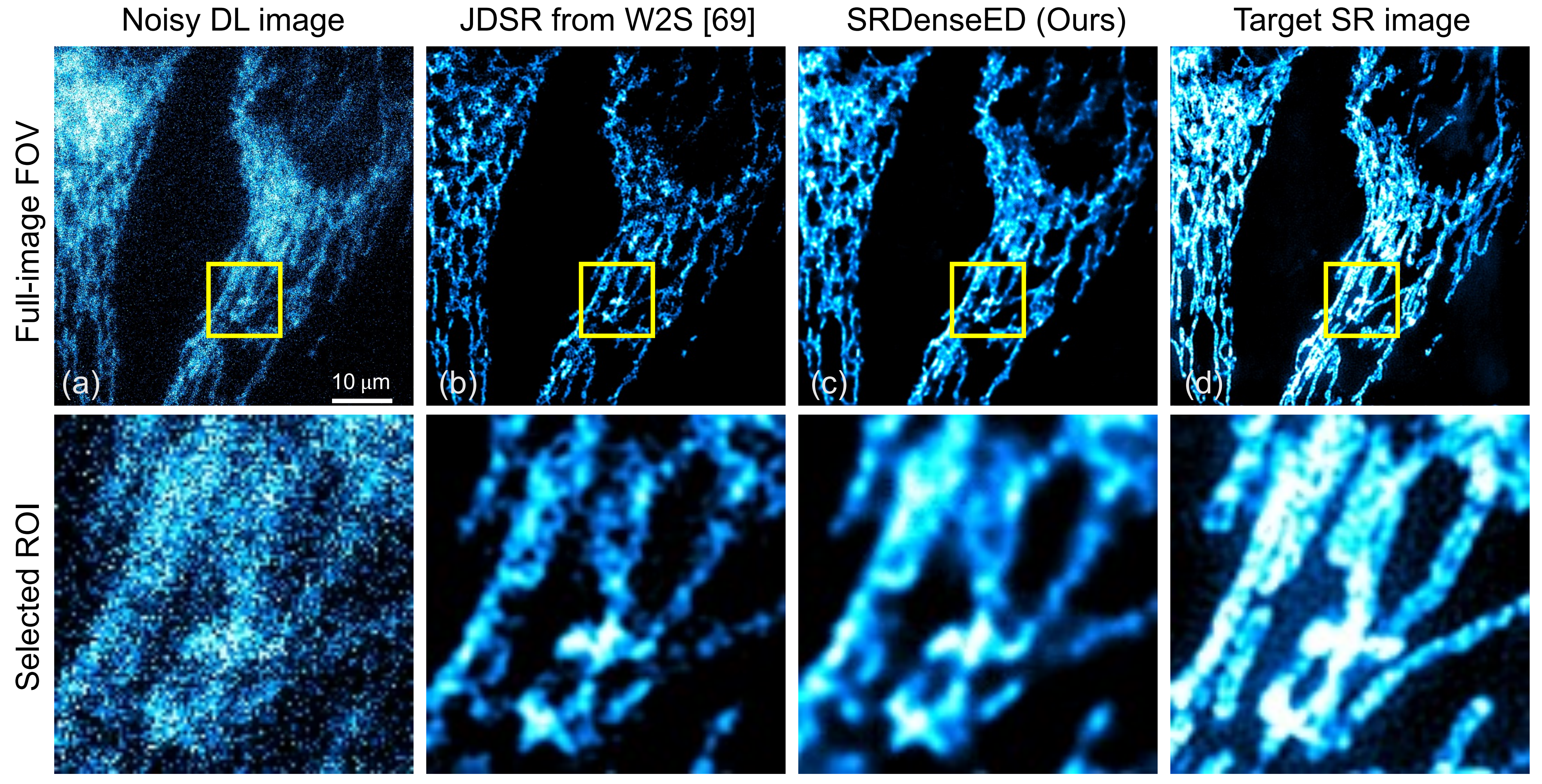

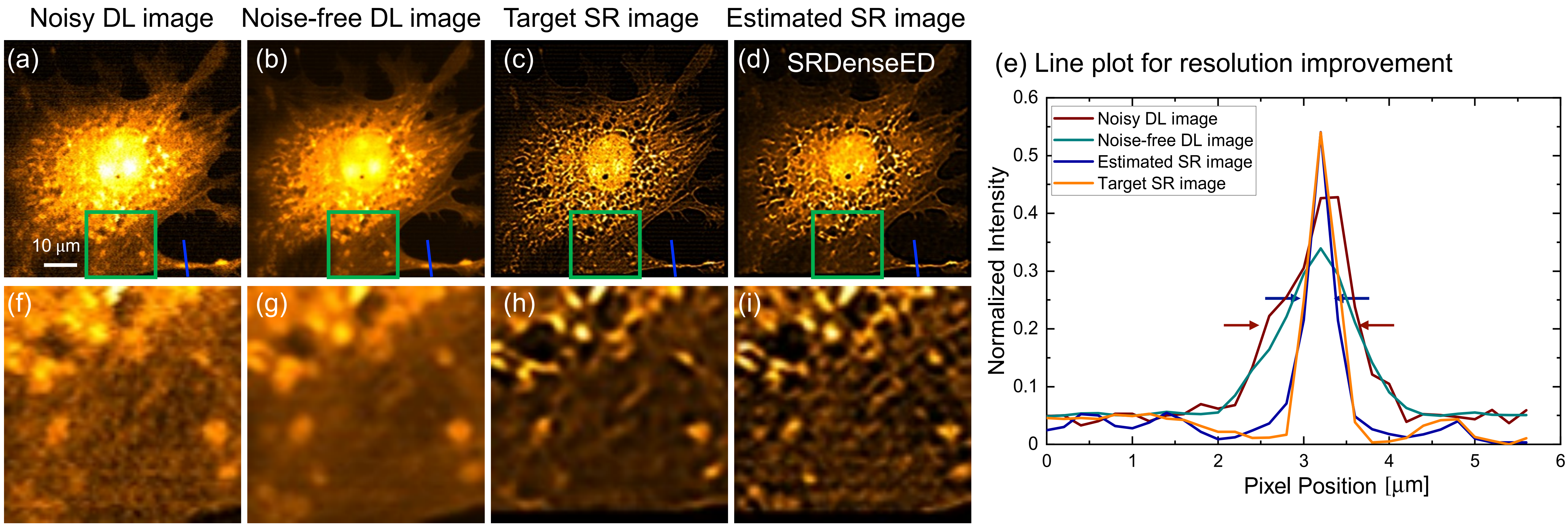
